# Genotype–Phenotype Distinctions in Spastic Paraplegia 4 Reveal HDAC6 as a Therapeutic Target

**DOI:** 10.1101/2025.07.15.664947

**Authors:** Neha Mohan, Skandha Ramakrishnan, Xiaohuan Sun, Ying Sun, Theresa Connors, Victor Chai, Emanuela Piermarini, Peter W. Baas, James Cai, Mei Liu, Liang Qiang

## Abstract

Spastic Paraplegia 4 (SPG4) is the most prevalent form of Hereditary Spastic Paraplegia (HSP), a neurodegenerative disorder characterized by progressive lower limb spasticity and debilitating gait impairment, primarily driven by axonal degeneration of corticospinal motor neurons (CSMNs). Caused by mutations in the *SPAST* gene encoding spastin, an AAA-ATPase involved in microtubule severing and intracellular organelle function, SPG4 accounts for 40–50% of autosomal dominant HSP cases, yet without effective treatments. Although reduced microtubule acetylation has emerged as a key pathological mechanism, whether and how distinct mutations lead to microtubule deacetylation and subsequent neurodegeneration remains unclear. To address this, we generated isogenic human induced pluripotent stem cell (hiPSC) lines with two distinct heterozygous *SPAST* mutations – SPAST^WT/C448Y^ (missense) and SPAST^WT/S245X^ (truncation). Employing an innovative differentiation protocol, we created human motor cortical organoids enriched in CSMNs, providing a robust platform to study SPG4 pathophysiology. These organoids revealed striking genotype-phenotype distinctions, with mutation-specific variations in CSMN loss, axonal degeneration and neuronal activities, mirroring clinical heterogeneity. Mechanistic studies identified aberrant activation of histone deacetylase 6 (HDAC6), a major neuronal microtubule deacetylase, as a key driver of SPG4 pathology. This dysregulation was specifically attributed to mutant M1-spastin, the longer isoform of spastin. Remarkably, pharmacological inhibition of HDAC6 with Tubastatin A restored microtubule acetylation status and mitigated axonal degeneration in both SPAST-mutant organoids, with corresponding improvements in corticospinal tract integrity and gait deficits validated in SPG4-transgenic mice. Collectively, our study establishes isogenic hiPSC-derived motor cortical organoids as a robust human model for corticospinal motor neuron degeneration and identifies HDAC6 hyperactivation as a central pathogenic mechanism and viable therapeutic target in SPG4.

## Introduction

Hereditary Spastic Paraplegia (HSP) is a genetically diverse neurodegenerative disorder characterized by progressive lower limb spasticity and weakness due to degeneration of the corticospinal tract (CST), the bundle of axons projected from corticospinal motor neurons (CSMNs) in the primary motor cortex. ^1–4^ Spastic Paraplegia 4 (SPG4), caused by mutations in *SPAST*, is the most prevalent autosomal dominant HSP subtype, accounting for ∼40% of genetically identified cases, with a prevalence of 2 to 6 per 100,000 individuals. ^5,6^ Clinically, SPG4 predominantly manifests as pure HSP, although complex forms of HSP exhibit additional features such as ataxia, cognitive impairment, and peripheral neuropathy. ^3,7–10^ Disease severity and age of onset vary substantially, yet the molecular bases underlying this genotype–phenotype variability remains poorly understood, hindering targetable therapy. ^11,12^ *SPAST* encodes spastin, an ATPases Associated with Diverse Cellular Activities (AAA) family protein essential for microtubule dynamics, intracellular organelle function and neuronal integrity. ^13–17^ Spastin has two isoforms: the longer ER-targeted M1-spastin, involved in ER morphology and membrane remodeling, and the shorter cytoplasmic M87-spastin, mainly responsible for microtubule severing. ^15,18–20^ While M87 is more abundant, accumulating evidence suggests mutant M1-spastin significantly drives SPG4 pathology through toxic gain-of-function mechanisms. ^21,22^ Our prior work suggested that this gain-of-function toxicity can sufficiently lead to neurodegeneration, with *SPAST* haploinsufficiency further exacerbating the pathology. ^23,24^ However, the isoform-specific contributions to each pathogenic mechanism remain undefined. ^20,25^ SPAST mutations, encompassing missense, truncation, frameshift, and splice-site variants, uniformly cause axonal degeneration but lead to highly variable clinical presentations. ^26^ Despite hypothesized roles for genetic modifiers and environmental factors, direct molecular insights into genotype–phenotype distinctions in SPG4 are missing. Emerging studies implicate microtubule hypoacetylation, driven by mutant M1-spastin accumulation, as a consistent pathological phenotype, regardless of the mutation. ^20,23,24,27^ Interestingly, histone deacetylase 6 (HDAC6), the primary cytoplasmic tubulin deacetylase in neurons, ^28–31^ is aberrantly activated in SPG4 transgenic models, though its pathological relevance in SPG4-patients remains unverified. ^24^ Indeed, HDAC6 inhibition has demonstrated therapeutic efficacy in many preclinical models of neurodegenerative diseases such as Alzheimer’s disease and Amyotrophic Lateral Sclerosis, its potential in the context of SPG4 has not been pursued. ^32–36^ Given the mechanistic involvement of HDAC6 in microtubule hypoacetylation and axonal degeneration, and the current lack of effective, targeted interventions for SPG4, systematically evaluating HDAC6 inhibition offers a compelling opportunity to address a critical unmet therapeutic need in this disorder.

To address these critical gaps, we established isogenic human induced pluripotent stem cell (hiPSC) lines carrying either a missense (SPAST^WT/C448Y^) or truncation (SPAST^WT/S245X^) mutation via CRISPR-Cas9 editing. These hiPSCs were differentiated into CSMN-enriched motor cortical organoids (MCOs), as the 3D human SPG4 model. Combined with validation in our established SPG4-transgenic mice, this novel platform enables precise interrogation of genotype–phenotype distinctions and aims to elucidate HDAC6 hyperactivity as a unifying mechanism and therapeutic target in SPG4.

## Materials and Methods

### Isogenic SPG4-hiPSC Generation, Validation and Cultures

Isogenic human induced pluripotent stem cells (hiPSCs) were derived from a control hiPSC line (FA0000010) which was generated from a healthy 36-year-old male donor using previously established protocols which was obtained from Columbia Stem Cell Initiative Stem Cell Core facility. ^37^ CRISPR (Clustered regularly interspaced short palindromic repeat) – Cas9 (CRISPR-associated protein 9) genome editing was employed to generate isogenic hiPSC lines harboring two distinct disease-associated mutations in the SPAST gene. Highly specific single-guide RNAs (sgRNAs) were designed to target defined loci within SPAST, enabling precise introduction of pathogenic variants via homology-directed repair, while preserving genomic integrity at off-target sites. All hiPSC lines and subcolonies were validated for their pluripotency by immunostaining for OCT4, SSEA-4, SOX2, and TRA1-60 (Thermo Fisher Scientific, A24881). Germ layer differentiation was conducted using the Human Pluripotent Stem Cell Functional Identification Kit (R&D Systems, SC027B) and G-banded karyotype and mutation analysis was also carried out to confirm CRISPR-Cas9 based mutation. All hiPSC lines were cultured in 5% CO_2_ at 37°C in mTeSR™ Plus (Stem Cell Technologies, 100-0276) on Matrigel™ GFR Basement Membrane Matrix (Corning, 356231) as previously reported. ^38^ Briefly, 6-well plate (Thermo Scientific, 140675) was coated with Matrigel diluted in ice-cold DMEM-F12 (Gibco, 11330032) for 1 hour at 37°C. Once all hiPSCs reach 70% confluence, they are passaged using ReLeSR™ (Stem Cell Technologies, 100-0483) which is incubated for 4 minutes and subsequently plated on coated dishes with 10μM ROCK inhibitor Y-27632 (Tocris, 1254). 3ml Medium changes with mTeSR™ Plus was performed every other day.

### Motor Cortical Organoid Development and Maintenance

Organoids were generated using a modified version of the previously reported publication. ^38,39^ Briefly, on day 0, pure and 90% confluent hiPSCs were lifted using ReLeSR for 4 minutes and plated onto ultra-low attachment 6-well plate (Corning, 3471) in mTeSR Plus with 10µM ROCK inhibitor Y-27632, 5µM Dorsomorphin (Sigma, C956T21), and 10µM SB431542 (Tocris, 1614) to promote embryoid bodies (EBs) formation. From day 3 to 6, a half medium changes with Dorsomorphin and SB431542 in mTeSR Plus was performed every other day. From days 7 to 23, half medium change was performed using neuronal differentiation media I which consist of Neurobasal A (Thermo Fisher, 10-888-022), B-27 without Vitamin A (Thermo Fisher, 12587010), Glutamax (Thermo Fisher, 35-050-061), 20ng/mL EGF (PeproTech, AF-100-15), 20ng/mL bFGF (Peprotech, 100-18B), particularly with 5µM cyclopamine (Fisher Scientific, AAJ61528MB). On day 23, organoids were placed on an orbital shaker (60 rpm) and the medium was switched to neuronal differentiation media II which consists of Neurobasal A (Thermo Fisher, 10-888-022), B-27 Plus (Thermo Fisher Scientific, A3582801), Glutamax (Thermo Fisher, 35-050-061), Normocin (InvivoGen, ant-nr-05), 20 ng/mL BDNF (PeproTech, 450-02) and 20 ng/mL NT3 (PeproTech, 450-03). All organoids were cultured in 5% CO2 at 37°C on an orbital shaker. Medium changes were performed every 3-4 days.

### Organoid Fixation, Section and Immunohistochemistry (IHC)

Organoids were washed once with PBS (Gibco, 10010023) and fixed in 4% PFA overnight at 4°C followed by dehydration in 30% sucrose until fully saturated. Organoids were then embedded in M1 Embedding Matrix (Epredia, 1310) and cryosectioned at 25µm thickness and placed on charged slides (Fisher Scientific, 1255016). The slides were dried at room temperature overnight. Organoid mounted slides were then hydrated and washed in 1X PBS 3 times for 5 mins followed by quenching (70% PBS, 30% methanol, 0.5% hydrogen peroxide) for 1 hour at room temperature. Subsequently, the slides were washed in PBS to remove the quenching buffer and incubated in either donkey (Jackson ImmunoResearch, 017-000-121) or goat (Jackson ImmunoResearch, 005-000-121) serum blocking buffer diluted in PBS for 1 hour at room temperature. Then primary antibodies listed in Supplementary Table 1 diluted in PBS and 0.1% Triton-X-100 was incubated overnight at 4°C followed by PBS washes and corresponding secondary antibodies conjugated to Alexa Fluor 488, 555, or 647 incubated for 2 hours at room temperature. Lastly, the slides were washed in PBS and incubated with DAPI at 1:20,000 dilution (Fisher Scientific, D1306) and mounted using Fluoro-Gel with Tris Buffer (Electron Microscopy Sciences 1798510). Organoid sections were imaged using Leica SP8 inverted confocal microscope and Leica Thunder microscope. Image J was used to measure average fluorescence intensity of each organoid section and individual neuronal processes.

### Single-cell RNA Sequencing (scRNA-seq)

10X Chromium Next GEM single cell 3’ Gene expression kit V3.1 dual index kit and 10X Chromium Controller system was used to generate single cell data. Briefly, 2.5-month-old organoids were dissociated using 4 ml StemPro Accutase™ Cell Dissociation Reagent (Thermo Fisher Scientific, A1110501) for 12 minutes at 37°C. Subsequently, the organoids were quenched and treated briefly with DNAase I (Thermo Fisher Scientific, EN0521) and gently triturated using 2ml serological pipettes to generate single cell suspension in PBS with 1% BSA. Cell viability and count were assessed using trypan blue staining and a hemocytometer. The cell suspension was then loaded onto Chromium next GEM Chip G (10x Genomics, 2000177) per manufacturer instructions, with a targeted cell recovery of 5000-10000 cells. Post GEM formation, the samples were subjected to reverse transcription, amplification, fragmentation and clean-ups, as recommended by the manufacturer. We then performed Illumina library preparation using the Dual Index plate TT Set A (10X Genomics, 3000431; Plate, 15535894). The subsequent library was then sequenced using NovaSeqxxx at a depth of 25000-30000 paired end reads per cell.

Raw sequencing data were initially processed and aligned to GRCh38 (human) using Cell Ranger Count v7.0.1 (10X Genomics, Cloud analysis) and HDF5 files were generated. The HDF5 files were then uploaded to the Partek Flow software with scRNAseq toolkit (Illumina). Using a single cell QA/QC and a noise-reduction pipeline, the cells were further filtered to exclude low quality cells and features. The filtered set was then normalized using the recommended counts per million, add 1, log 2 transformation workflow. Principal component analysis (PCA) was performed on the filtered cells and further clustered using Louvain graph-based clustering algorithm using the top 20 PCAs and visualized using uniform manifold approximation and projection (UMAP). We then extracted the biomarkers for these clusters and used them to manually annotate the clusters. Panglao database, Human protein atlas and extensive literature review were used to annotate the seven clusters as neuroepithelial cells, glial cells, intermediate progenitor cells, immature corticospinal motor neurons, deep layer excitatory neurons, GABAergic neurons and glutamatergic neurons. A heat map was generated using the marker genes of each cell cluster.

### RNA Isolation and qRT-PCR

RNA was isolated from various organoid samples using the PureLink™ RNA Mini Kit (Thermo Fisher Scientific, 12183018A) and subsequently reverse transcription was performed using the High-Capacity cDNA Reverse Transcription Kit (Thermo Fisher Scientific, 4368813) to synthesize cDNA. Quality control and quantification was performed using the NanoDrop Spectrophotometer. qRT-PCR was performed using the 2X Universal SYBR Green Fast qPCR Mix (ABclonal, RK21203) on the StepOnePlus™ Real-Time PCR System (Applied Biosystems). RNA primers were acquired from Integrated DNA Technologies and listed in Supplementary Table 1.

### Multielectrode Array (MEA) Recording and Analysis

Single-wells containing 60 electrodes (Multi Channel Systems, MEA2100-Systems), were coated with Poly-L-Ornithine (Sigma Aldrich, P4957) overnight at room temperature then washed 3-5 times with sterile ddH_2_O. The wells were fully dried, and organoids were placed centrally covering all the electrodes and cultured in BrainPhys Neuronal Medium (Stem Cell Technologies, 05790) supplemented with BDNF and NT3. Two days after organoid placement, extracellular spontaneous neuronal activity was recorded for a duration of 5 mins using the Multi-Channel Experimenter software as previously described. ^40^ The recording parameter was acquired at 20 kHz and filtered with a 3500 Hz fourth-order high-pass Butterworth filter and neuronal spikes were detected when the signals surpassed a threshold of 6 standard deviations from the baseline noise. Bursts were detected when ≥ 4 spikes with a duration of 50 milliseconds (ms) and an interval of 100ms between bursts occurred. Network bursts were detected when a minimum of 10 electrodes were active with 5 of them simultaneously participating. Acquired data was processed using Multi Channel Analyzer and organized in Microsoft Excel.

### SPG4-dHet Transgenic Mouse Colony Maintenance

All the transgenic mouse experiments were performed in compliance with the NIH’s Guide for the Care and Use of Laboratory Animals and were reviewed and approved by the Institutional Animal Care and Use Committee at Drexel University. Five mice were housed per cage under a 12-hour light/dark cycle, temperature and humidity were kept constant, and mice had free access to drinking water and food. Colony generation, breeding strategies, and genotyping are all previously described in detail. ^23,24^ For this study, we used our novel double-heterozygous mice (dHet) that contains one copy of human mutant SPAST^C448Y^ at Rosa26 locus and one copy of endogenous mouse *Spast* (hSPAST-C448Y^+/-^; mSpast^-/+^) which depicts both CST dieback and adult-onset gait deficiencies that are remarkably reminiscent of human patients. ^24^

### Preparation and Systemic Delivery of Tubastatin A in Mice

Tubastatin A (Selleckchem, S8049, Batch: S804913) was prepared according to manufacturer’s instructions. Animals were administrated Tub A in 5% DMSO and 95% corn oil at a dose of 16mg/kg with a dosing volume of 10ml/kg daily for 21 days intraperitoneally. A mixture of DMSO and corn oil was used as a vehicle. After 21 days of treatment, animals were euthanized for biochemical analysis. For organoid studies, 100µM Tub A in DMSO was added to the culture media daily for 72 hours and collected for downstream analysis.

### Catwalk Assay

The Noldus Catwalk XT is an automated gait analysis system for assessing locomotion in rats and mice. The behavioral platform and procedure were previously described in great detail. ^23,24^ Briefly, mice were isolated in the behavior room 30 mins prior to testing to acclimate them to the new environment. Each mouse underwent a week of training where they learned the task following a week of recording for the various gait parameter analysis. Animals were individually placed on the catwalk to freely move in both directions. After 15 mins, the animal was removed from the walkway and returned to its home cage. The camera gain was set to 20dB and the detection threshold was set to 0.10. Compliant runs were classified as run with a duration between 0.50 and 5.00 seconds with a maximum speed variation of 60%. For each animal, 5-6 compliant runs per group were used for analysis. For consistency and rigor, both training and recording were conducted by the same person. Five to 6 animals per group were used for behavioral studies. Clinically relevant parameters were chosen for analysis for this study. ^24,41^

### Spinal Cord Tissue Processing and Quantitative Axonal Analysis

Spinal cord tissue collection, processing, and anatomical analysis were performed as previously described. ^23,24^ For anatomical analysis in this study, three mice per group were sacrificed by intraperitoneal injection of 150 mg/kg Euthasol solution (VEDCO, 50989056912). Transcardial perfusion was performed using a buffered 0.9% NaCl rinse followed by tissue fixation in a mixture of 4% paraformaldehyde (PFA; Electron Microscopy Sciences, 19202) and 1% Glutaraldehyde in 0.1M phosphate buffer. Spinal cords were dissected out and were post-fixed overnight at 4°C in the same fixative solution. Cords were then washed in 0.1M phosphate buffer overnight at 4°C and cervical and lumbar blocks of spinal cord were sectioned at 200µm using a Vibratome. Sectioned cords were further post-fixed in cold buffered 2% osmium tetroxide for 1 hour followed by sequential dehydration in 70, 95 and 100% ethanol followed by two rinses in propylene oxide (PO). The cords were then incubated in a 1:1 mixture of Epon -Araldite and PO for 1 hour followed by a 2:1 mixture of Epon-Araldite and PO overnight and lastly incubated in 100% Epon-Araldite with 2% DMP-30 hardener for 2 hours. The spinal cord sections were finally embedded in fresh Epon-Araldite with 2% DMP-30 hardener in silicone molds and allowed to polymerize for 72 hours in a 60°C oven. 1µm sections were cut from each block using a glass knife mounted on a ultramicrotome and sections were stained with toluidine blue to visualize cross-sections of axons. The corticospinal tract located in the most ventral part of the dorsal column was imaged using Axio Observer 7 Zeiss microscope with a 100X objective. Axonal count and perimeter were quantified using ImageJ.

### Whole Blood Collection and Serum Preparation

Whole blood was collected from 3 mice per group. Whole blood was collected through a transcardial puncture and processed in Vacutainer® Serum Tubes (Becton Dickinson, 366668) following a previously published protocol. ^42^ To isolate the serum from whole blood, the blood was incubated at room temperature for 15 mins to clot and centrifuged for 10 mins at 4°C at 3000g. The resulting supernatant was collected and sent to the Animal Diagnostic Laboratory at the University of Michigan Medical School-Pathology Core for biochemical analysis to assess hepatotoxicity.

### HDAC6 Activity Analysis

HDAC6 activity assays were performed according to the manufacturer’s instructions (BPS Biosciences, 50076-1) and as previously described. ^23,24^ Briefly, treated and untreated organoid samples, as well as spinal cord tissue samples, were lysed in HDCA6 lysis buffer and centrifuged at 16,000g for 10 mins at 4°C. The protein content resulting from the supernatant was quantified using the Pierce BCA Protein Assay Kit (Thermo Fisher Scientific, 23227). Samples were then incubated at 37°C for 30 mins with HDAC6 substrate followed by incubating the developer for 10 mins at 37°C to stop the reaction. The fluorescence was measured on a Tecan spectral plate reader at 380/490nm (excitation/emission). The analysis was conducted per the manufacturer’s instructions where the results are expressed as U per mg of protein, where U stands for the amount of HDAC6 required to deacetylate 1pmol of HDAC6 substrate per min.

### Spastin Isoform Overexpression in SH-SY5Y Cells

SH-SY5Y neuroblastoma cell line was used to overexpress either wild-type or mutant forms of M1 or M87 as listed in Supplementary Table 1. Cells were cultured in DMEM/F12 medium (Thermo Fisher Scientific, 11320033) supplemented with 10% fetal bovine serum (Novus Biologicals, S11150) and 1% penicillin-streptomycin (Pen/Strep, 100 IU/ml and 100 µg/ml), and plated in 100mm culture dishes (BD Falcon, 353003). At 60-70% confluency, cells were transfected with 2µg of wildtype or mutant M1/M87 constructs using Lipofectamine 2000 (Invitrogen, 11668-019) according to the manufacturer’s instructions. 72 hours post transfection, cells were harvested in ice-cold PBS and centrifuged at 1500 rpm x 5 minutes at room temperature. The resulting cell pellets were further lysed and used for HDAC6 activity assays (see above). Non-transfected (naïve) SH-SY5Y cells were included as control.

### Protein Extraction and Western Blotting

Organoid samples were lysed in 1X RIPA Buffer (Thermo Fisher Scientific, 89900) supplemented with Halt™ Protease and Phosphatase Inhibitor Cocktail (Thermo Fisher Scientific, 78446). For animal studies, the mice are euthanized as described above and the motor cortex and spinal cords were dissected and homogenized in the same RIPA buffer with inhibitors. All samples were sonicated on ice with two 30-second on/off pulses, then centrifuged at 15,000 × g for 30 minutes at 4 °C. The supernatant was collected and total protein concentration determined using the Pierce BCA Protein Assay Kit (Thermo Fisher Scientific, 23227). Equal amounts of protein (25 µg per sample) from both organoid and mouse tissue lysates were combined with 4X LDS sample buffer (Thermo Fisher Scientific, B0007) and 10X Sample Reducing Agent (Thermo Fisher Scientific, B0009), and then subjected to SDS-PAGE on 4–12% Bis-Tris Plus gels (Thermo Fisher Scientific, NW04120BOX). Proteins were transferred onto PVDF membranes (Thermo Fisher Scientific, 88518) using either a 0.34 A constant current for 2 hours (instant transfer) or 0.1 A overnight on ice. Following transfer, membranes were blocked at room temperature for 1 hour in LI-COR blocking buffer (Li-co Bioscience, 927-60010), then incubated overnight at 4°C with primary antibodies as listed in Supplementary Table 1. The next day, membranes were incubated with fluorescent secondary antibodies (LI-COR IRDye 680RD or IRDye^®^ 800CW)) for 2 hours at room temperature on a shaker. Protein bands were visualized using the Odyssey^®^ DLx Infrared Imaging System (LI-COR), and band intensities quantified using Fiji (ImageJ).

### Co-immunoprecipitation (Co-IP) and Western Blotting

Co-immunoprecipitation was performed to investigate potential interactions between HDAC6 and spastin isoforms. Due to the lack of suitable antibodies for immunoprecipitating endogenous spastin and HDAC6, epitope-tagged expression constructs were employed. HEK293T cells were co-transfected with mcherry-HDAC6 and one of the following Tet-on, FLAG-tagged spastin constructs: control vector (CON255-FLAG), M1-Spastin^WT^-FLAG, M87-Spastin^WT^-FLAG, M1-Spastin^C448Y^-FLAG, M1-Spastin^S245X^-FLAG as listed in Supplementary Table 1. The CON255-FLAG plasmid was included as a negative control. After 48 hours of induction with doxycycline (5 µg/mL), total protein was extracted using IP lysis buffer (Thermo Fisher Scientific, 87787) supplemented with protease (NCM Biotech, P001) and phosphatase inhibitors (NCM Biotech, P003). For FLAG immunoprecipitation, 500 µL of lysate was incubated with 50 µL of anti-FLAG magnetic agarose beads (Thermo Fisher Scientific, A36797) at room temperature for 20 minutes with continuous rotation. Beads were then washed three times with PBS to remove nonspecific proteins. Bound complexes were eluted by boiling the beads in 50 µL of 1× SDS loading buffer for 5 minutes, followed by SDS-PAGE and western blot analysis. For reverse Co-IP, lysates were processed similarly but incubated with RFP-Trap agarose beads (NanoTag Biotechnologies, N0410) to immunoprecipitate mcherry-HDAC6 and examine potential association with spastin isoforms. Eluates were resolved and analyzed by western blotting as described above. Briefly, 5μg input control samples and 15μL IP protein samples were separated using SDS-PAGE and transferred to PVDF membrane. The membrane was blocked at room temperature for 2 hours in 5% non-fat dried milk (BioFroxx, 1172GR500) in Tris-buffered saline (TBS, pH 7.4). The samples were then incubated with anti-Flag (1: 2000, Proteintech, 20543-1-AP), anti-Phosphoserine/threonine (1:1000, Amyjet Scientific, PPS-PP2551), anti-HDAC6 (1:1000, Affinity Biosciences, AF6485) or anti-mCherry (1:2000, Abcam, ab205402) at 4°C overnight as listed in Supplementary Table 1. After washing with TBST (TBS with 0.1% Tween 20), goat anti-Rabbit-IgG (1:2000, Jackson ImmunoResarch Laboratories) or goat anti-chicken-IgG (1:2000, Jackson ImmunoResarch Laboratories) antibodies were applied at room temperature for 2 hours. The blot was covered with ECL solution (Tanon, 180-5001) and visualized with Chemiluminescence imaging system (Tanon).

### Statistical Analysis and Data Visualization

All control and experimental groups were processed in parallel and analyzed blindly to minimize variability and enhance reproducibility. Data were organized in Microsoft Excel and statistical analysis and graph preparations were carried out using GraphPad Prism version 10. All data sets were subjected to the Shapiro-Wilk’s test for normality and outliers were eliminated using the “Identify outliers” program in GraphPad Prism version 10. Statistical comparisons were performed using one-way ANOVA followed by Tukey’s post hoc test or unpaired t-tests, with p < 0.05 considered statistically significant. Figures, graphs, and schematics were assembled using Adobe Photoshop 2024.

## Results

### Derivation of CSMN-enriched MCOs from isogenic hiPSC lines carrying distinct SPAST mutations

To rigorously investigate genotype–phenotype relationships and faithfully recapitulate SPG4 pathology, we generated isogenic hiPSC lines harboring either a missense (SPAST^WT/C448Y^) or a truncation mutation (SPAST^WT/S245X^), both introduced heterozygously into a single parental hiPSC line derived from a healthy 36-year-old male donor, accurately reflecting the autosomal dominant inheritance pattern of SPG4. These lines exhibited normal colony morphology, pluripotency expression (Figure 1A), and differentiation capability into all three germ layers (Figure 1B), with confirmed stable male karyotype (Figure 1C). The precise introduction of SPAST mutations via CRISPR-Cas9 was validated by targeted PCR and sequencing for both SPAST^WT/S245X^ (Figure 1D) and SPAST^WT/C448Y^ (Figure 1E). MCOs enriched with CSMNs were generated from all three hiPSCs using our optimized protocol, adapted from previously established methods (Figure 1F). ^38,39^ Briefly, embryoid bodies (EBs) derived from hiPSCs underwent neuronal induction with dual SMAD inhibition, followed by neuronal patterning in the presence of growth factors and the sonic hedgehog (SHH) inhibitor cyclopamine, which is essential for enriching the CSMN population ^43,44^ followed by neuronal maturation. 1-month-old motor cortical organoids from all three hiPSC lines showed robust populations of neural progenitors marked by Sox2 and postmitotic neurons marked by tau and MAP2 (Figure 1G-H’) with significant neuroepithelial loops resembling active neurogenesis (Figure 1H’). Note that no overt difference was identified among the lines. At 3 months of differentiation, MCOs exhibited substantial enrichment of CSMNs, as confirmed by prominent expression of established CSMN-specific markers including CTIP2, L1CAM, and vGlut1 (Figures 1I–M’). Importantly, 3-month-old organoids also demonstrated progressive neuronal maturation, characterized by extensive expression of mature neuronal marker, NeuN, and the axonal marker, phosphorylated neurofilament, SMI312, robust populations of astrocytes exhibiting typical astrocytic morphology and immunoreactivity for GFAP were observed, indicating the establishment of complex neuronal-glial interactions (Figures 1K-M’). By 6 months of differentiation, MCOs displayed dense networks of synapses, as evidenced by widespread expression of synaptophysin (Figure 1N-N’). To further validate functional neuronal activity, we utilized a multielectrode array (MEA) system, which confirmed spontaneous neuronal firing and network activity within organoids (Figures 1O-Q). Additionally, single-cell RNA sequencing (scRNA-seq) was performed on 2.5-month-old healthy hiPSC-derived organoids, resulting in the identification of 4,819 cells expressing a total of 25,052 genes (Figure 1R). Unbiased clustering and manual annotation delineated seven distinct cell populations characterized by unique marker gene expression profiles (Figures 1S–T, Supplementary file). These clusters closely resemble cell populations found in the developing human motor cortex, underscoring the fidelity and physiological relevance of our organoid model. This further highlights the validity and region-specificity of our MCO model. Quantitative qRT-PCR analysis provided further molecular validation of CSMN identity. Compared to conventional forebrain cortical organoids generated without cyclopamine treatment, our motor cortical organoids demonstrated significantly higher expression of the canonical CSMN markers, including CTIP2, L1CAM, CRYM and vGlut1 (Figures 1U–Y, see Supplementary Table 2 for mean ± SD). ^45^ Despite prior reports of UCHL1 labeling CSMNs in mice ^46,47^, unlike other CSMN markers, our data reveal no difference between pan-cortical and motor cortical organoids, aligning with its broad neuronal expression. ^48,49^

**Figure 1.**
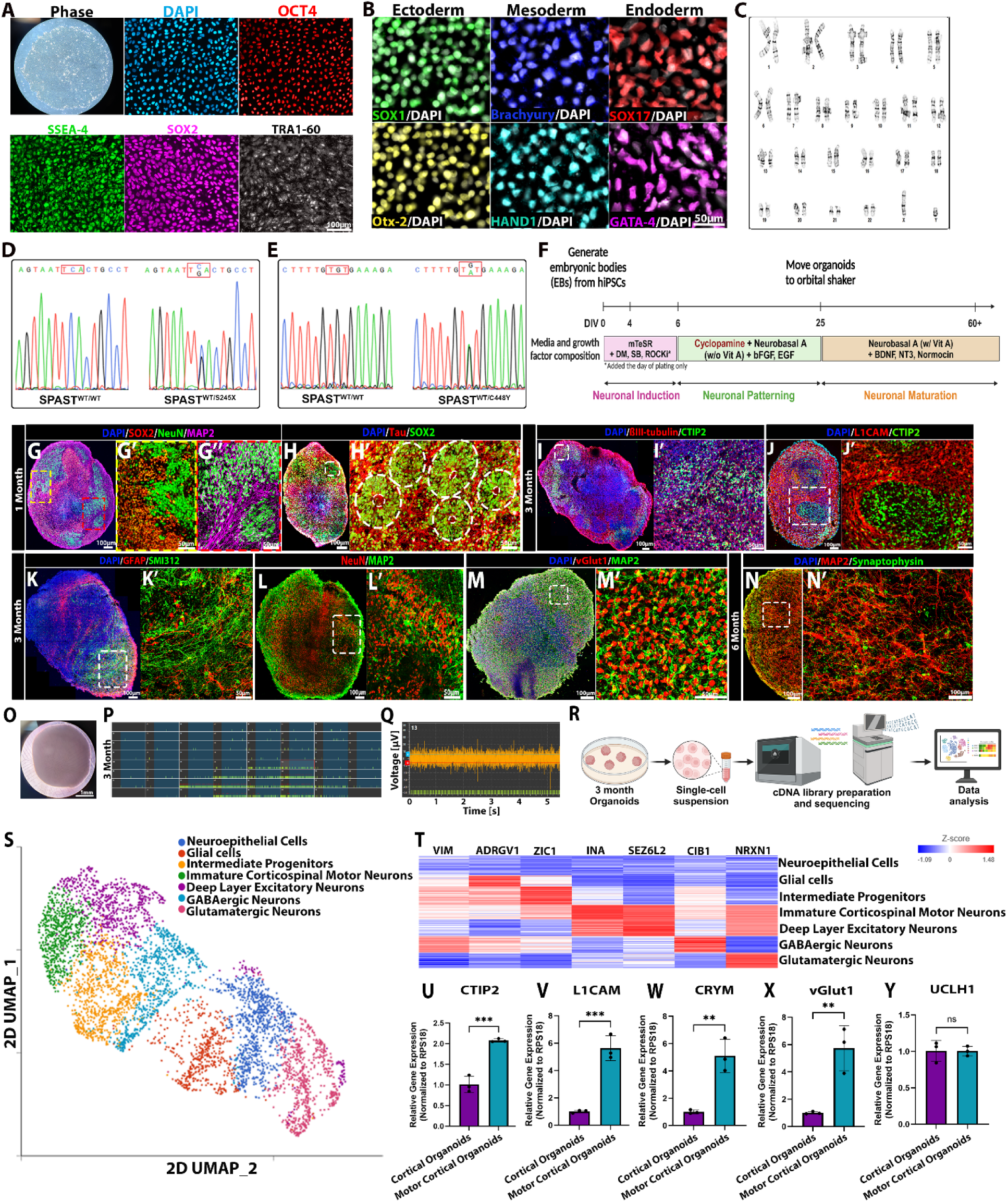
Derivation and characterization of CSMN-enriched MCOs from isogenic hiPSC lines carrying distinct SPAST mutations. A. Representative phase-contrast and immunofluorescent images of hiPSC colonies (SPAST^WT/WT^) including DAPI and pluripotent markers OCT4, SSEA-4, SOX2, and TRA-60. B. Representative immunofluorescent images for tri-germ layer differentiation of isogenic hiPSCs (SPAST^WT/WT^), including SOX17 and GATA4 (endoderm), Brachyury and HAND1 (mesoderm), and SOX1 and Otx2 (ectoderm). C. Normal karyotypes were identified across all three isogenic hiPSC lines, represented by SPAST^WT/WT^. D-E. Genomic DNA sequencing confirming the specific genotypes of hiPSC-SPAST^WT/S245X^, hiPSC-SPAST^WT/C448Y^ and hiPSC-SPAST^WT/WT^. F. Schematic and timeline of MCO generation. G-H’. Representative immunofluorescent images of 1-month MCOs using pan-neuronal (NeuN, Tau, MAP2) and neural progenitor (SOX2) markers. Dashed circles highlight neuroepithelial loops. I-M’. Representative immunofluorescent images of 3-month MCOs using pan-neuronal (βIII-tubulin, SMI312), astrocytic (GFAP) and CSMN (CTIP2, L1CAM, vGlut1) markers. N-N’. Representative immunofluorescent images of 6-month MCOs using presynaptic marker (synaptophysin). O. Representative phase image of a 3-month MCO docked on a MEA2100 single-well platform. P. Representative image showing electrode setup and spontaneous neuronal firing – green lines indicate spike counts, red bars indicate bursts, and blue highlight indicates network burst. Q. Raster plot from P showing individual spikes from electrode number 13 with a standard deviation of ± 6 from baseline recording (+6: blue line and −6: red line). R. Schematic of scRNA-seq workflow. S. UMAP of scRNAseq of 2.5-month-old MCOs. T. Heatmap of S with cell-type specific marker genes (see supplementary file for details). U-Y. qRT-PCR analysis of CSMN specific genes, normalized to RSP18 in 6-month generic cortical organoids (n=3) vs. MCOs (n=3). Unpaired t-test. **p<0.05, ***p<0.001. All data shown as mean ± SD, see Supplementary Table 1 for details.

### Distinct genotype-phenotype distinctions in isogenic SPG4 organoids highlight accelerated axonal degeneration and selective CSMN vulnerability in the missense mutation model

SPG4 presents with marked clinical heterogeneity, with mutation type strongly influencing disease onset and progression, suggesting underlying genotype–phenotype distinction. ^50^ To investigate these differences in our model, we first examined spastin mRNA expression in 6-month-old MCOs using qRT-PCR. Primers targeting both the N-terminal (exons 1–2) and C-terminal (exons 11–12) regions revealed elevated full-length spastin transcripts in both SPAST^WT/S245X^ and SPAST^WT/C448Y^ organoids compared to wild-type controls (Figures 2A–B). This was particularly surprising for SPAST^WT/S245X^, which might be expected to undergo nonsense-mediated mRNA decay yet aligns with previous patient data showing preserved or elevated SPAST mRNA levels despite truncating mutations. ^51^ No significant difference was observed between the two mutant groups at the transcript level. At the protein level, SPAST^WT/S245X^ organoids showed modest reduction in full-length M1- and M87-spastin, whereas SPAST^WT/C448Y^ organoids displayed increased levels of both isoforms (Figures 2C-E). Notably, a truncated M1-spastin species was detected in the SPAST^WT/S245X^ organoids, reinforcing the possibility that aberrant accumulation of mutant protein may confer gain-of-function toxicity in SPG4 truncation mutants.

**Figure 2.**
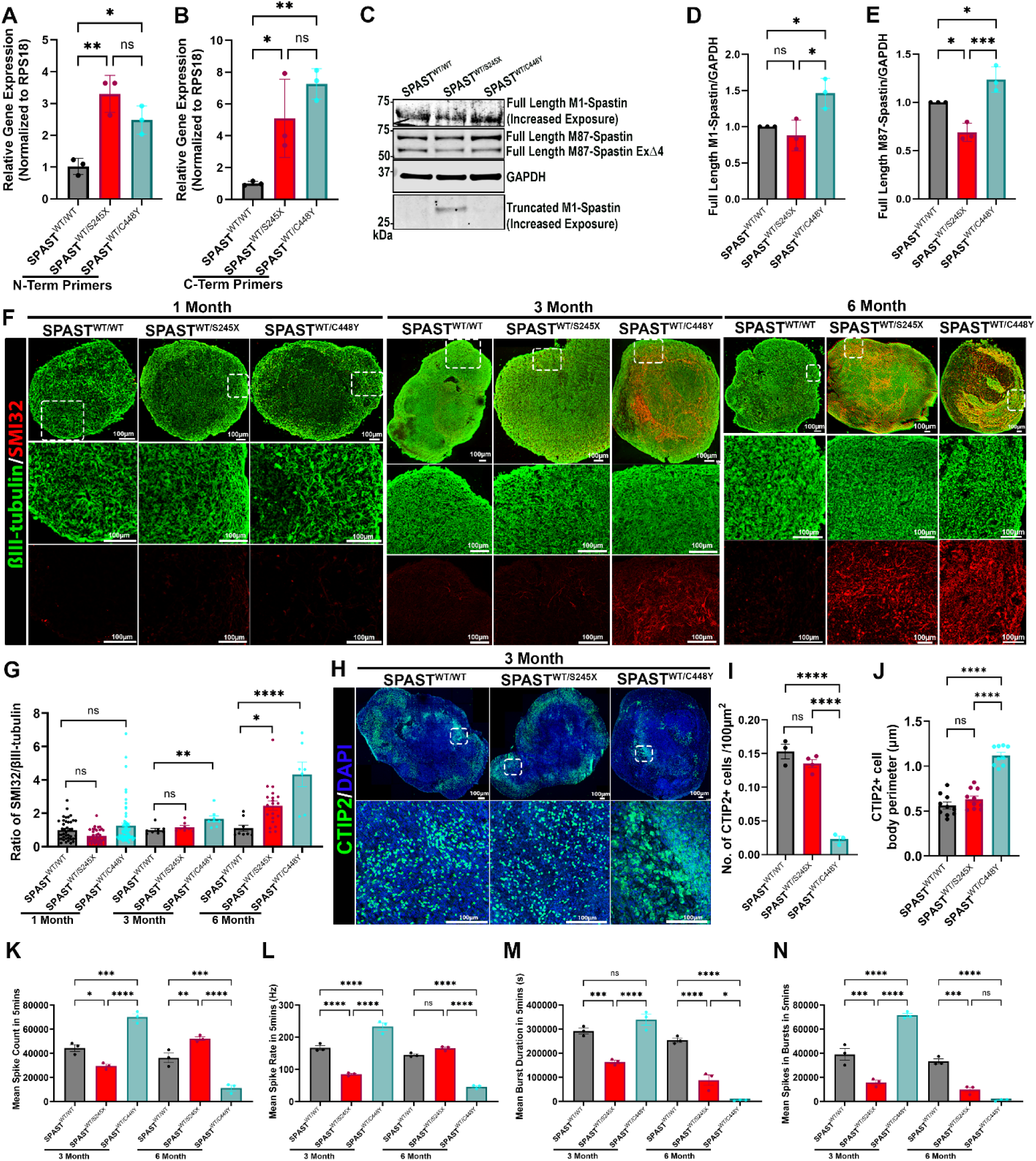
Genotype-phenotype distinctions in isogenic SPG4 MCOs highlight accelerated axonal degeneration and selective CSMN vulnerability in SPAST^WT/C448Y^. A-B. Characterization of spastin expression in 6-month MCOs derived from the three isogenic genotypes (n=3 organoids), normalized to RSP18 revealed by qRT-PCR with N-terminal primers (A) and C-terminal primers (B). C-E. Representative western Blot (WB) of spastin in 6-month-old MC organoids (C, n=3). Levels of full length M1-spastin (D) and M87-spastin (E) in SPAST^WT/S245X^ and SPAST^WT/C448Y^, both normalized to GAPDH relative to SPAST^WT/WT^. Note that a truncated product of SPAST^WT/S245X^ MCOs has been identified with predicted molecular weight. F-G. Representative immunofluorescent images of 1-, 3-, and 6-month MCOs from all three genotypes marked by SMI32, the early axonal degeneration marker (F) normalized to βIII-tubulin and quantified (G). H-J. Representative immunofluorescent images of CSMN populations marked by CTIP2 in 3-month MCOs from all three genotypes (H, n=3) and the quantification analyses of the numbers (I) and cell body perimeters (J) of the CTIP2^+^ cells. K-N. Spontaneous extracellular neuronal activity over 5-minute recording of 3-, and 6-month MCOs from all three genotypes (n=3). The graphs show mean spike count that passed the threshold (K), mean spike rate (L), mean burst duration that surpasses a minimum of 50 milliseconds (ms) (M), and a mean number of spikes to be considered a burst (N). One-way ANOVA with Tukey *post hoc* analysis. *p<0.05, **p<0.005, ***p<0.001. ****p<0.0001. All data shown as mean ± SD, see Supplementary Table 2 for details.

Given prior clinical findings linking AAA domain missense mutations to earlier onset and more severe SPG4 phenotypes ^26,50,52^ than the truncation mutations, we next sought to determine whether our model recapitulates these differential degenerative trajectories. We performed a temporal analysis of axonal degeneration using SMI32, a dephosphorylated neurofilament marker of axonal degeneration. ^53,54^ At 1 month, neither mutant showed signs of axonal degeneration. However, by 3 months, SPAST^WT/C448Y^ organoids exhibited a significant increase in SMI32 signal, which intensified by 6 months (Figure 2F). In contrast, SPAST^WT/S245X^ organoids only showed enhanced SMI32 at the 6-month time point (Figures 2F-G), suggesting a delayed degenerative course. Corroborating the axonal degeneration phenotype, we observed a marked reduction in CSMNs, identified by the expression of CTIP2, in 3-month-old SPAST^WT/C448Y^ organoids compared to isogenic wild-type controls (Figures 2H-I). This selective neuronal loss was accompanied by a significant enlargement of CTIP2-positive soma, consistent with cytoskeletal dysregulation and early signs of neurodegeneration (Figure 2J) ^55^. Notably, these phenotypes – including CSMN loss and somatic hypertrophy – were absent in SPAST^WT/S245X^ organoids, further underscoring the mutation-specific nature of the SPG4 neurodegenerative progression. To further dissect the genotype-phenotype differences, we employed electrophysiological recordings using single-well MEA platform to record extracellular neuronal activity on mature MCOs. 3-month-old SPAST^WT/C448Y^ MCOs displayed neuronal hyperactivity denoted by increased spike count and rate (Figures 2K-L) along with increased burst duration and the number of spikes in bursts (Figures 2M-N) whereas the SPAST^WT/S245X^ MCOs demonstrated reduced neuronal activity. Subsequently, at 6 months, SPAST^WT/C448Y^ MCOs exhibit diminished neuronal activity with the SPAST^WT/S245X^ MCO following suit for burst duration and number of spikes in bursts however spike count and rate show neuronal hyperactivity (Figure 2K-N, see Supplementary Table 3 for mean ± SD). This bi-phasic change in neuronal activity coincides with progressive axonal degeneration. These data reveal that our novel motor cortical organoids recapitulate key genotype-phenotype features of SPG4, including the earlier and more severe neuronal pathology associated with AAA-domain missense mutations.

### Differential yet convergent HDAC6 hyperactivation induced microtubule hypoacetylation underlies the axonal degeneration in distinct *SPAST*-mutant organoids

Given the distinct yet partially overlapping neurodegenerative phenotypes observed in SPAST^WT/C448Y^ and SPAST^WT/S245X^ MCOs, we next investigated whether these mutation-specific outcomes converge on shared downstream mechanisms. We focused on dysregulation of microtubule acetylation – previously implicated as a key pathological feature in SPG4 models. ^23,24^ While aberrant HDAC6 activation induced microtubule hypoacetylation was identified in one SPG4 mouse model, ^24^ its relevance across diverse SPAST mutations remains unclear. Thus, we examined microtubule acetylation and the corresponding HDAC6 activity at different time points in our mutant organoids. At 3 months, SPAST^WT/C448Y^ organoids exhibited a marked reduction in microtubule acetylation and elevated HDAC6 activity, while SPAST^WT/S245X^ organoids retained near-normal levels of both (Figures 3A–B, G). By 6 months, however, both mutant lines displayed significant microtubule hypoacetylation and HDAC6 hyperactivation, further highlighting delayed pathological onset in SPAST^WT/S245X^ (Figures 3A, C, H) Importantly, these changes in HDAC6 activity occurred without alterations in total HDAC6 protein levels (Figures 3I–J), suggesting possible posttranslational dysregulation. This temporal divergence in HDAC6-microtubule defects mirrors the genotype-specific pattern of axonal degeneration, reinforcing a mechanistic link between HDAC6 activation and neurodegenerative progression in SPG4.

**Figure 3.**
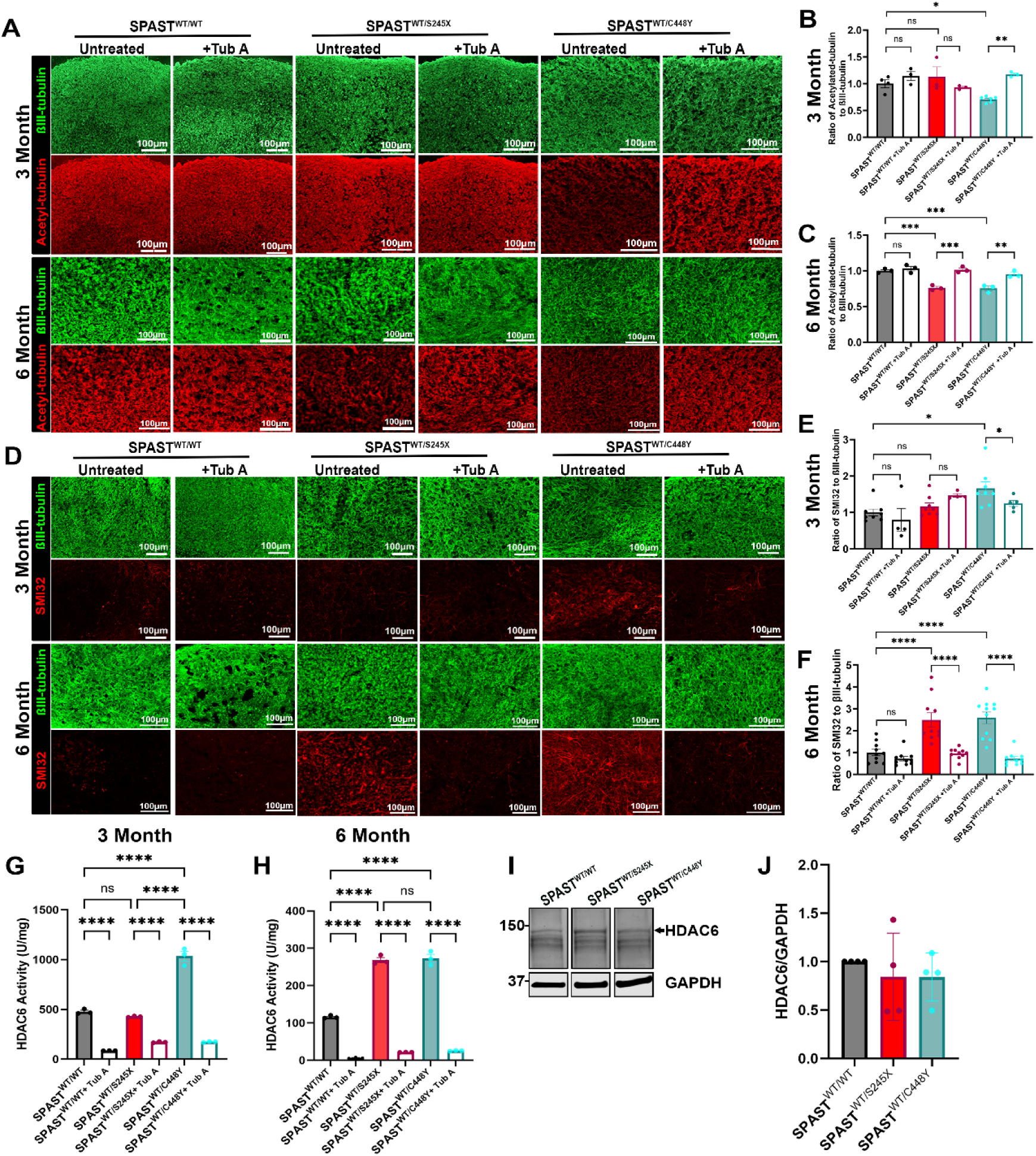
HDAC6 hyperactivation induced by mutant M1-spastin drives microtubule hypoacetylation and axonal degeneration in SPAST-mutant MCOs and represents a promising therapeutic target. A. Representative immunofluorescent images of 3- and 6-month MCOs derived from SPAST^WT/WT^, SPAST^WT/S245X^ and SPAST^WT/C448Y^ lines, untreated or treated with the HDAC6 inhibitor Tub A, stained for acetylated tubulin and βIII-tubulin. B-C. Quantification of the acetylated tubulin to βIII-tubulin ratio in 3-month (B) and 6-month (C) MCOs (n=3-6). Tub A significantly increased tubulin acetylation in mutant MCOs. D. Representative immunofluorescent images of 3- and 6-month-old MCOs from SPAST^WT/WT^, SPAST^WT/S245X^ and SPAST^WT/C448Y^ lines, untreated or treated with TubA, stained for SMI32 (a marker of early axonal degeneration) and βIII-tubulin. E-F. Quantification of the SMI32 to βIII-tubulin ratio in individual axons from 3-month (n=8, E) and 6-month (n=10, F) MCOs. Tub A treatment significantly reduced SMI32 levels in mutant organoids. G-H. Quantification of HDAC6 enzymatic activity in 3-month (n=3, G) and 6-month (n=3, H) MCOs from SPAST^WT/WT^, SPAST^WT/S245X^ and SPAST^WT/C448Y^ lines, with or without Tub A treatment. HDAC6 activity is expressed as units (U) per mg of total protein, where one unit is defined as the amount of enzyme required to deacetylate 1pmol of a synthetic acetylated peptide substrate per minute. Elevated HDAC6 activity in mutant organoids is significantly reduced by Tub A treatment at both time points. I-J. WB (I) and quantification (J) of HDAC6 (black arrow) protein levels in 3-month MCOs derived from SPAST^WT/WT^, SPAST^WT/S245X^ and SPAST^WT/C448Y^ lines. HDAC6 expression, normalized to GAPDH and shown relative to SPAST^WT/WT^, revealed no significant differences across genotypes (n = 4). One-way ANOVA with Tukey *post hoc* analysis. *p<0.05, **p<0.002, ***p<0.001, ****p<0.0001. All data shown as mean ± SD, see Supplementary Tables 3-4 for details.

To evaluate the therapeutic potential of targeting HDAC6, we treated organoids with Tubastatin A (Tub A), a selective HDAC6 inhibitor. We evaluated the optimal dose of Tub A treatment in organoids in a dose dependent manner (data not shown) in conjunction with previously published studies. ^56–58^ 100µM Tub A treatment successfully suppressed HDAC6 hyperactivity and normalized microtubule acetylation in both genotypes at their respective stages of pathology (Figures 3A–C, G–H). Interestingly, Tub A significantly reduced SMI32 accumulation and improved axonal morphology, indicating structural rescue of degeneration (Figures 3D–F, see Supplementary Tables 4-5 for mean ± SD). These findings establish HDAC6 hyperactivation and microtubule hypoacetylation as temporally distinct yet converging mechanisms in SPG4 pathogenesis, driven by mutation-specific dynamics. Moreover, they validate HDAC6 inhibition as a mechanistically grounded, genotype-spanning therapeutic strategy.

### HDAC6 selectively interacts with M1-spastin and is aberrantly activated by mutant forms

Given that both SPAST^WT/S245X^ and SPAST^WT/C448Y^ organoids exhibit HDAC6 hyperactivation and microtubule hypoacetylation in addition to our previous findings that mutant M1-spastin rather than M87-spastin, irrespective of mutation type, consistently drives this phenotype. Next, we sought to determine whether these effects arise from a direct, isoform-specific interaction between spastin and HDAC6. Co-immunoprecipitation (co-IP) assays in HEK293T cells expressing mCherry-tagged HDAC6 and Flag-tagged spastin isoforms revealed robust interaction between HDAC6 and all M1 variants (WT, C448Y, S245X), whereas no interaction was observed with wild-type M87-spastin (Figures 4A–B). Additionally, co-IP examination using anti-Flag beads further confirmed this M1-specific association with endogenous HDAC6 (Figures 4C-D). Quantitative analyses revealed that both M1-SPAST^S245X^ and M1-SPAST^C448Y^ exhibited increased HDAC6 binding relative to wild-type, with M1-SPAST^C448Y^ showing the highest level of interaction (Figure 4E). Moreover, HDAC6 co-immunoprecipitated with M1-SPAST^C448Y^ displayed significantly elevated phosphorylation, as detected by a phospho-HDAC6–specific antibody, suggesting enhanced enzymatic activation compared to wild-type and truncation variants (Figure 4C). To determine whether HDAC6 hyperactivation is a shared downstream effector of mutant spastin toxicity and whether this effect is isoform-specific, we quantified HDAC6 enzymatic activity in SH-SY5Y cells expressing wild-type or mutant M1- or M87-spastin. ^22^ HDAC6 activity remained unaltered in cells expressing either wild-type M1-spastin or any form of M87-spastin. In contrast, all mutant M1-spastins markedly increased HDAC6 activity, regardless of mutation type (Figure 4E, see Supplementary Tables 6-7 for mean ± SD). These findings implicate mutant M1-spastin, irrespective of mutation type, as a key driver of HDAC6 hyperactivation and support a convergent pathological mechanism underlying SPG4.

**Figure 4.**
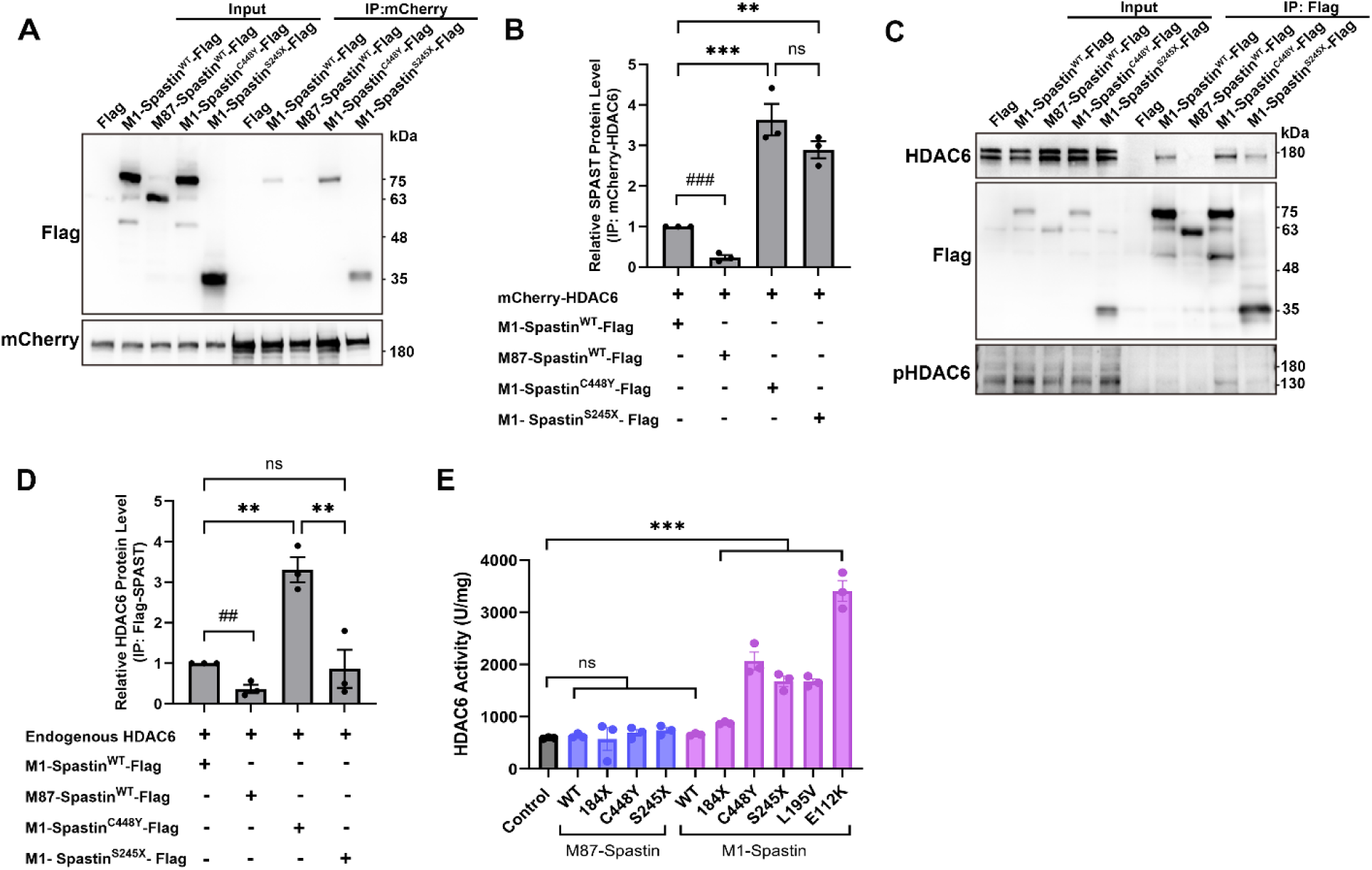
HDAC6 selectively interacts with M1-spastin and is aberrantly activated by mutant forms. A-B. Co-immunoprecipitation (Co-IP) in HEK293T cells co-expressing HDAC6-mCherry and Flag-tagged spastin isoforms revealed that HDAC6 binds selectively to M1-spastin, but not M87-spastin (A). Quantification confirmed robust enrichment of M1-spastin isoforms, in the HDAC6-containing immunoprecipitated complex, with significantly increased interaction observed for mutant forms (B). C-D. Reciprocal Co-IP using anti-Flag antibody was performed to isolate spastin isoforms, followed by WB with anti-HDAC6 and anti-phospho-HDAC6 antibodies, showing M1-Spastin^C448Y^-Flag immunoprecipitated more phosphorylated HDAC6. (C). Quantification also confirmed that mutant M1-spastin isoforms pulled down greater amounts of HDAC6 compared to controls (D). E. HDAC6 activity assays in SH-SY5Y cells expressing various M1 and M87 spastin mutants showed that only M1 mutations led to significantly elevated HDAC6 enzymatic activity, indicating an isoform-specific gain-of-function effect. One-way ANOVA with Tukey *post hoc* analysis. **p<0.002, ***p<0.001, and Unpaired t-test. ##p<0.005, ###p<0.001. All data shown as mean ± SD, see Supplementary Tables 5-6 for details.

### Pharmacological inhibition of HDAC6 ameliorates gait deficiency and CST degeneration in SPG4-transgenic mice

To validate our organoid-based findings *in vivo*, we employed a novel double heterozygous (dHet) SPG4-transgenic mouse model (hSPAST-C448Y^+/-^; m*Spast*^-/+^), which harbors one allele of the SPAST^C448Y^ mutation – the same to our missense mutant hiPSC line – and one null allele of endogenous mouse *Spast*. This recently developed mouse model recapitulates some key aspects of human SPG4 by combining gain-of-function toxicity and haploinsufficiency, exhibiting pathological phenotypes including progressive gait impairment, CST degeneration, and axonal swelling. ^24^ Consistent with our organoid data, dHet mice also exhibit reduced microtubule acetylation and aberrant HDAC6 hyperactivation. ^24^ We treated 3-month-old male dHet mice – corresponding to symptom onset – with Tub A via daily intraperitoneal injection for 21 days (Figure 5A). Serum chemistry confirmed no hepatotoxicity, indicating favorable tolerability (Supplementary Table 8). Gait analysis using the CatWalk XT system revealed significant hindlimb deficits in dHet mice, including decreased print width, length, area, and altered paw placement – phenotypes reminiscent of SPG4 patient presentations – which were significantly ameliorated following Tub A treatment (Figures 5B–F). As expected, no forelimb differences were detected, in line with previous observations. ^24^ Furthermore, Tub A effectively suppressed hyperactivated HDAC6 and restored microtubule acetylation without altering overall neuronal microtubule mass (Figures 5G–J). Anatomical evaluation of CST axons via toluidine blue staining of semi-thin spinal cord sections revealed reduced axonal density and increased swelling in the ventral portion of the lumbar dorsal column of dHet mice, as previous reported, ^24^ which were partially rescued by Tub A treatment (Figures 5K–N, see Supplementary Table 9 for mean ± SD). No significant changes were detected at the cervical level. Together, these *in-vivo* results substantiate HDAC6 hyperactivation as a disease-relevant, therapeutically modifiable mechanism in SPG4. The convergence between organoid and animal model findings strongly supports selective HDAC6 inhibition as a viable strategy for mitigating CST degeneration and motor deficits in SPG4 patients.

**Figure 5.**
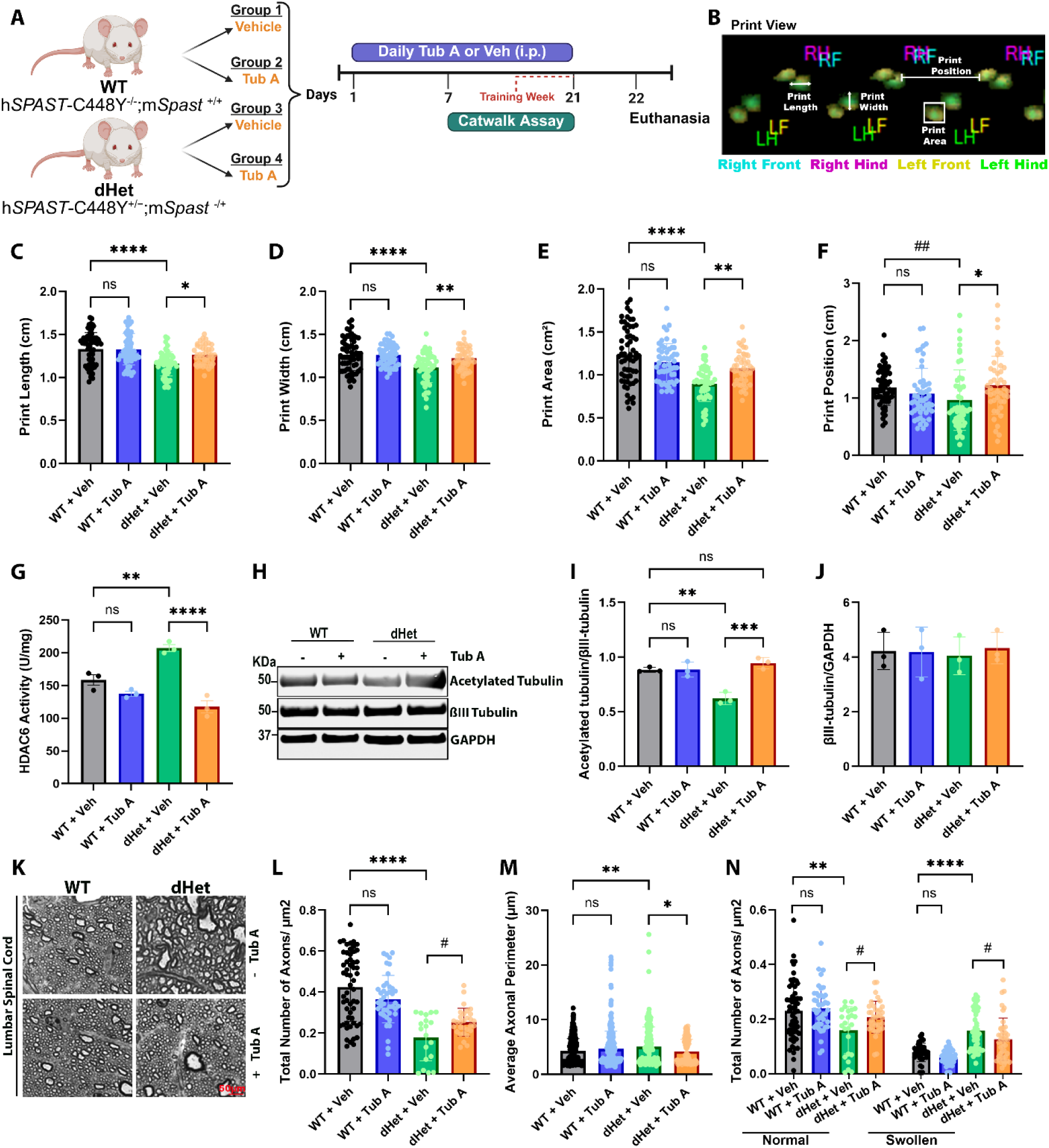
HDAC6 inhibition with TubA improves the gait deficiency and restores CST integrity in dHet mice. A. Experimental timeline of daily Tub A or vehicle treatment and CatWalk analysis in WT and dHet mice. B. Representative CatWalk print view showing paw positioning and parameters analyzed, including print length, width, area, and position. C–F. Quantitative gait metrics across experimental groups: Print length (C), Print width (D), Print area (E), and Print position (F) (hind paw placement relative to the ipsilateral front paw). Data represent classified runs from 6 animals per group. G. HDAC6 enzymatic activity measured from spinal cord lysates across groups (n=3 mice per group). H-J. WB analysis of lumbar spinal cord lysates showing levels of acetylated tubulin and βIII-tubulin in WT and dHet mice treated with Tub A or vehicle. Tub A treatment significantly increases acetylated tubulin levels in dHet mice. βIII-tubulin levels remain unchanged across groups and serve as a loading control (n=3 per group). K. Representative toluidine blue-stained cross-sections of the dorsal column in lumbar spinal cords from treated and untreated WT and dHet mice. L. Quantitative analyses of CST axons: total axon count per μm² (L), average axonal perimeter (M), and number of morphologically normal versus swollen axons (based on shape classification as normal: regular, swollen: irregular (N). Axons with a perimeter of ≥ 10µm was considered swollen). Data represents cumulative axon counts (n=120-500) from 3 mice per group. One-way ANOVA with Tukey *post hoc* analysis. *p<0.05, **p<0.002, ***p<0.001, ****p<0.0001 and Unpaired t-test. #p<0.05, ##p<0.005. All data shown as mean ± SD, see Supplementary Table 8 for details.

## Discussion

Our study establishes an innovative, physiologically relevant platform for modeling SPG4 using CSMN-enriched MCOs derived from isogenic hiPSCs. Unlike generic cortical organoids, our MCOs uniquely replicate cellular populations specific to the motor cortex, particularly CSMNs, the primary neuronal subtype affected in SPG4. ^59–61^ Utilizing an isogenic background, this system enables precise interrogation of genotype–phenotype correlations for missense (SPAST^WT/C448Y^) and truncation (SPAST^WT/S245X^) mutations, addressing the previously elusive molecular underpinnings of clinical variability observed in SPG4. ^50^ Clinically, missense *SPAST* mutations generally cause earlier onset and more severe phenotypes compared to truncation mutations. Our findings align closely with these clinical discoveries, revealing accelerated and selective CSMN degeneration in SPAST^WT/C448Y^ organoids. Mechanistically, elevated *SPAST* mRNA levels were observed in both mutant organoids, confirming previous reports that *SPAST* mutations can escape nonsense-mediated decay. ^51^ Notably, accumulation of both M1- and M87-spastin isoforms specifically in SPAST^WT/C448Y^ organoids indicates that missense mutations may confer a toxic gain-of-function component – an effect inherently absent in *Spast*-knockout models that only reflect haploinsufficiency pathology. ^62,63^ The detection of truncated spastin proteins in SPAST^WT/S245X^ organoids, raises the possibility that truncation mutations may also exert toxic gain-of-function effects, rather than acting solely through haploinsufficiency. ^22^ Furthermore, our co-IP studies revealed that while both M1-SPAST^S245X^ and M1-SPAST^C448Y^ isoforms interact with HDAC6, the missense mutant M1-SPAST^C448Y^ exhibits markedly stronger binding, correlating with greater HDAC6 hyperactivation. This differential interaction suggests that distinct *SPAST* mutations may drive varying degrees of HDAC6 dysregulation, contributing to mutation-specific pathogenic severity in SPG4 – a pattern that is recapitulated in our isogenic organoids with time-course studies. Indeed, including more patient-derived hiPSC lines and additional *SPAST* mutations in our organoid models will be important for fully capturing SPG4’s genetic and phenotypic diversity and advancing mechanistic and therapeutic insights.

Our analysis using both MCOs and transgenic mice revealed microtubule hypoacetylation as a key pathological feature in SPG4. This defect was significantly more pronounced in the missense mutant and closely correlated with greater neurodegenerative severity in SPG4 patients. ^50,62^ A prior study using a single, non-isogenic hiPSC line harboring an intronic splice-site *SPAST* mutation reported increased microtubule acetylation in 2D-cultured neurons, a result that differs from our findings. ^64^ Notably, this outcome phenocopies findings from spastin-depletion models, suggesting a phenotype driven predominantly by the haploinsufficiency mechanism in that context. ^24,65,66^ In contrast, our isogenic organoid models and a larger patient-derived cohort study, ^67^ both report microtubule hypoacetylation, supporting a more consistent pathogenic signature across SPG4 mutations with better-controlled experimental systems. Indeed, the contrasting findings also underscore the molecular heterogeneity of SPG4 and emphasize the need to disentangle distinct genotype–phenotype relationships, particularly in models that incorporate disease-relevant cell types and contexts. Last but not the least, our findings highlight HDAC6 hyperactivation, driven by mutant M1-spastin, as a convergent and therapeutically relevant mechanism in SPG4 pathogenesis. The ability of Tubastatin A to restore microtubule acetylation and ameliorate neurodegenerative phenotypes across both organoid and animal models underscores the translational potential of HDAC6 inhibition, offering a new avenue for therapeutic intervention for SPG4. Note that the partial rescue achieved with HDAC6 inhibition in adult mice highlights promising avenues for future studies, including exploring combination therapies that could enhance therapeutic efficacy, especially in advanced stages of disease progression.

Taken together, our study unveils CSMN-enriched MCOs as a groundbreaking platform to decode the elusive corticospinal neuron vulnerability and advancing accelerating targeted therapy development for SPG4 and other CSMN-related neurodegenerative diseases.

## Acknowledgements

We thank Angela Altadonna for assisting in breeding mice for the experiments and Larisa Ibric and Luisa Abadia for assisting with behavioral studies. We would like to thank the staff of the Columbia Stem Cell Initiative Stem Cell Core facility, under the leadership of Dr. Barbara Corneo at Columbia University Irving Medical Center for assisting the CRISPR editing for the work, including Dr. Grazia Iannello, Dr. Dario Sirabella, Dr. Achchhe Patel, Dr. Hemanta Sarmah and Ms. Ekaterina Lebayle. We also want to thank Dr. Joanne Mathiasen for providing robust discussion and insights into dosing Tubastatin A in mice as well as Dr. Matthias Kneussel for providing the spastin knockout mice for breeding to generate the dHet mouse. All schematics were generated using BioRender.

## Supplemental Materials

Supplementary file: spreadsheet file containing details regarding marker genes of identified cell populations for scRNA-seq, related to Figure 1.

Supplementary Table 1: List of antibodies, primers, and plasmids used for this study.

Supplementary Table 2: Mean ± SD values from Figure 1.

Supplementary Table 3: Mean ± SD values from Figure 2.

Supplementary Table 4: Mean ± SD values from Figure 3.

Supplementary Table 5: Mean ± SD values from Figure 3.

Supplementary Table 6: Mean ± SD values from Figure 4.

Supplementary Table 7: Mean ± SD values from Figure 4.

Supplementary Table 8: Chemistry panel of serum extracted from whole blood samples to evaluate hepatotoxicity with Tub A treatment, related to Figure 5.

Supplementary Table 9: Mean ± SD values from Figure 5.

## Data Availability

The datasets used and analyzed during the current study are available from the corresponding author upon request.

## References

1. Solowska, J.M., and Baas, P.W. (2015). Hereditary spastic paraplegia SPG4: what is known and not known about the disease. Brain 138, 2471–2484. 10.1093/brain/awv178.

2. Shribman, S., Reid, E., Crosby, A.H., Houlden, H., and Warner, T.T. (2019). Hereditary spastic paraplegia: from diagnosis to emerging therapeutic approaches. Lancet Neurol 18, 1136–1146. 10.1016/s1474-4422(19)30235-2.

3. Blackstone, C. (2018). Converging cellular themes for the hereditary spastic paraplegias. Curr Opin Neurobiol 51, 139–146. 10.1016/j.conb.2018.04.025.

4. Reid, E. (2003). Science in motion: common molecular pathological themes emerge in the hereditary spastic paraplegias. J Med Genet 40, 81–86. 10.1136/jmg.40.2.81.

5. Ruano, L., Melo, C., Silva, M.C., and Coutinho, P. (2014). The global epidemiology of hereditary ataxia and spastic paraplegia: a systematic review of prevalence studies. Neuroepidemiology 42, 174–183. 10.1159/000358801.

6. Mackay-Sim, A. (2021). Hereditary Spastic Paraplegia: From Genes, Cells and Networks to Novel Pathways for Drug Discovery. Brain Sci 11. 10.3390/brainsci11030403.

7. Bruyn, R.P. (1992). The neuropathology of hereditary spastic paraparesis. Clin Neurol Neurosurg 94 Suppl, S16–18. 10.1016/0303-8467(92)90010-z.

8. Fink, J.K. (2014). Hereditary spastic paraplegia: clinical principles and genetic advances. Semin Neurol 34, 293–305. 10.1055/s-0034-1386767.

9. Montanaro, D., Vavla, M., Frijia, F., Aghakhanyan, G., Baratto, A., Coi, A., Stefan, C., Girardi, G., Paparella, G., De Cori, S., et al. (2020). Multimodal MRI Longitudinal Assessment of White and Gray Matter in Different SPG Types of Hereditary Spastic Paraparesis. Front Neurosci 14, 325. 10.3389/fnins.2020.00325.

10. White, K.D., Ince, P.G., Lusher, M., Lindsey, J., Cookson, M., Bashir, R., Shaw, P.J., and Bushby, K.M. (2000). Clinical and pathologic findings in hereditary spastic paraparesis with spastin mutation. Neurology 55, 89–94. 10.1212/wnl.55.1.89.

11. Lo Giudice, T., Lombardi, F., Santorelli, F.M., Kawarai, T., and Orlacchio, A. (2014). Hereditary spastic paraplegia: clinical-genetic characteristics and evolving molecular mechanisms. Exp Neurol 261, 518–539. 10.1016/j.expneurol.2014.06.011.

12. Salinas, S., Proukakis, C., Crosby, A., and Warner, T.T. (2008). Hereditary spastic paraplegia: clinical features and pathogenetic mechanisms. Lancet Neurol 7, 1127–1138. 10.1016/s1474-4422(08)70258-8.

13. Kuo, Y.W., Trottier, O., Mahamdeh, M., and Howard, J. (2019). Spastin is a dual-function enzyme that severs microtubules and promotes their regrowth to increase the number and mass of microtubules. Proc Natl Acad Sci U S A 116, 5533–5541. 10.1073/pnas.1818824116.

14. Yu, W., Qiang, L., Solowska, J.M., Karabay, A., Korulu, S., and Baas, P.W. (2008). The microtubule-severing proteins spastin and katanin participate differently in the formation of axonal branches. Mol Biol Cell 19, 1485–1498. 10.1091/mbc.e07-09-0878.

15. Ji, Z., Zhang, G., Chen, L., Li, J., Yang, Y., Cha, C., Zhang, J., Lin, H., and Guo, G. (2018). Spastin Interacts with CRMP5 to Promote Neurite Outgrowth by Controlling the Microtubule Dynamics. Dev Neurobiol 78, 1191–1205. 10.1002/dneu.22640.

16. Park, S.H., Zhu, P.P., Parker, R.L., and Blackstone, C. (2010). Hereditary spastic paraplegia proteins REEP1, spastin, and atlastin-1 coordinate microtubule interactions with the tubular ER network. J Clin Invest 120, 1097–1110. 10.1172/jci40979.

17. Wang, X., and O’Kane, C.J. (2008). Hereditary spastic paraplegia genes in Drosophila: dissecting their roles in axonal degeneration and intracellular traffic. SEB Exp Biol Ser 60, 161–182.

18. Papadopoulos, C., Orso, G., Mancuso, G., Herholz, M., Gumeni, S., Tadepalle, N., Jüngst, C., Tzschichholz, A., Schauss, A., Höning, S., et al. (2015). Spastin binds to lipid droplets and affects lipid metabolism. PLoS Genet 11, e1005149. 10.1371/journal.pgen.1005149.

19. Connell, J.W., Allison, R.J., Rodger, C.E., Pearson, G., Zlamalova, E., and Reid, E. (2020). ESCRT-III-associated proteins and spastin inhibit protrudin-dependent polarised membrane traffic. Cell Mol Life Sci 77, 2641–2658. 10.1007/s00018-019-03313-z.

20. Costa, A.C., and Sousa, M.M. (2022). The Role of Spastin in Axon Biology. Front Cell Dev Biol 10, 934522. 10.3389/fcell.2022.934522.

21. Solowska, J.M., D’Rozario, M., Jean, D.C., Davidson, M.W., Marenda, D.R., and Baas, P.W. (2014). Pathogenic mutation of spastin has gain-of-function effects on microtubule dynamics. J Neurosci 34, 1856–1867. 10.1523/jneurosci.3309-13.2014.

22. Solowska, J.M., Rao, A.N., and Baas, P.W. (2017). Truncating mutations of SPAST associated with hereditary spastic paraplegia indicate greater accumulation and toxicity of the M1 isoform of spastin. Mol Biol Cell 28, 1728–1737. 10.1091/mbc.E17-01-0047.

23. Qiang, L., Piermarini, E., Muralidharan, H., Yu, W., Leo, L., Hennessy, L.E., Fernandes, S., Connors, T., Yates, P.L., Swift, M., et al. (2019). Hereditary spastic paraplegia: gain-of-function mechanisms revealed by new transgenic mouse. Hum Mol Genet 28, 1136–1152. 10.1093/hmg/ddy419.

24. Piermarini, E., Akarsu, S., Connors, T., Kneussel, M., Lane, M.A., Morfini, G., Karabay, A., Baas, P.W., and Qiang, L. (2022). Modeling gain-of-function and loss-of-function components of SPAST-based hereditary spastic paraplegia using transgenic mice. Hum Mol Genet 31, 1844–1859. 10.1093/hmg/ddab367.

25. Ramakrishnan, S., Mohan, N., Dong, Z., Liu, M., and Qiang, L. (2025). Unraveling Isoform Complexity: The Roles of M1- and M87-Spastin in Spastic Paraplegia 4 (SPG4). Mov Disord 40, 420–430. 10.1002/mds.30072.

26. Fonknechten, N., Mavel, D., Byrne, P., Davoine, C.S., Cruaud, C., Bönsch, D., Samson, D., Coutinho, P., Hutchinson, M., McMonagle, P., et al. (2000). Spectrum of SPG4 mutations in autosomal dominant spastic paraplegia. Hum Mol Genet 9, 637–644. 10.1093/hmg/9.4.637.

27. Leo, L., Weissmann, C., Burns, M., Kang, M., Song, Y., Qiang, L., Brady, S.T., Baas, P.W., and Morfini, G. (2017). Mutant spastin proteins promote deficits in axonal transport through an isoform-specific mechanism involving casein kinase 2 activation. Hum Mol Genet 26, 2321–2334. 10.1093/hmg/ddx125.

28. Osseni, A., Ravel-Chapuis, A., Thomas, J.L., Gache, V., Schaeffer, L., and Jasmin, B.J. (2020). HDAC6 regulates microtubule stability and clustering of AChRs at neuromuscular junctions. J Cell Biol 219. 10.1083/jcb.201901099.

29. Zhang, Y., Kwon, S., Yamaguchi, T., Cubizolles, F., Rousseaux, S., Kneissel, M., Cao, C., Li, N., Cheng, H.L., Chua, K., et al. (2008). Mice lacking histone deacetylase 6 have hyperacetylated tubulin but are viable and develop normally. Mol Cell Biol 28, 1688–1701. 10.1128/mcb.01154-06.

30. Zhang, Y., Li, N., Caron, C., Matthias, G., Hess, D., Khochbin, S., and Matthias, P. (2003). HDAC-6 interacts with and deacetylates tubulin and microtubules in vivo. Embo j 22, 1168–1179. 10.1093/emboj/cdg115.

31. Kalebic, N., Sorrentino, S., Perlas, E., Bolasco, G., Martinez, C., and Heppenstall, P.A. (2013). αTAT1 is the major α-tubulin acetyltransferase in mice. Nat Commun 4, 1962. 10.1038/ncomms2962.

32. Trzeciakiewicz, H., Ajit, D., Tseng, J.H., Chen, Y., Ajit, A., Tabassum, Z., Lobrovich, R., Peterson, C., Riddick, N.V., Itano, M.S., et al. (2020). An HDAC6-dependent surveillance mechanism suppresses tau-mediated neurodegeneration and cognitive decline. Nat Commun 11, 5522. 10.1038/s41467-020-19317-4.

33. Fazal, R., Boeynaems, S., Swijsen, A., De Decker, M., Fumagalli, L., Moisse, M., Vanneste, J., Guo, W., Boon, R., Vercruysse, T., et al. (2021). HDAC6 inhibition restores TDP-43 pathology and axonal transport defects in human motor neurons with TARDBP mutations. Embo j 40, e106177. 10.15252/embj.2020106177.

34. Osseni, A., Ravel-Chapuis, A., Belotti, E., Scionti, I., Gangloff, Y.G., Moncollin, V., Mazelin, L., Mounier, R., Leblanc, P., Jasmin, B.J., and Schaeffer, L. (2022). Pharmacological inhibition of HDAC6 improves muscle phenotypes in dystrophin-deficient mice by downregulating TGF-β via Smad3 acetylation. Nat Commun 13, 7108. 10.1038/s41467-022-34831-3.

35. d’Ydewalle, C., Krishnan, J., Chiheb, D.M., Van Damme, P., Irobi, J., Kozikowski, A.P., Vanden Berghe, P., Timmerman, V., Robberecht, W., and Van Den Bosch, L. (2011). HDAC6 inhibitors reverse axonal loss in a mouse model of mutant HSPB1-induced Charcot-Marie-Tooth disease. Nat Med 17, 968–974. 10.1038/nm.2396.

36. Simões-Pires, C., Zwick, V., Nurisso, A., Schenker, E., Carrupt, P.A., and Cuendet, M. (2013). HDAC6 as a target for neurodegenerative diseases: what makes it different from the other HDACs? Mol Neurodegener 8, 7. 10.1186/1750-1326-8-7.

37. Patel, A., Garcia Diaz, A., Moore, J.C., Sirabella, D., and Corneo, B. (2020). Establishment and characterization of two iPSC lines derived from healthy controls. Stem Cell Res 47, 101926. 10.1016/j.scr.2020.101926.

38. Sun, X., Kofman, S., Ogbolu, V.C., Karch, C.M., Ibric, L., and Qiang, L. (2024). Vascularized Brain Assembloids With Enhanced Cellular Complexity Provide Insights Into the Cellular Deficits of Tauopathy. Stem Cells 42, 107–115. 10.1093/stmcls/sxad086.

39. Yates, P.L., Case, K., Sun, X., Sullivan, K., Baas, P.W., and Qiang, L. (2022). Veteran-derived cerebral organoids display multifaceted pathological defects in studies on Gulf War Illness. Front Cell Neurosci 16, 979652. 10.3389/fncel.2022.979652.

40. Yates, P.L., Patil, A., Sun, X., Niceforo, A., Gill, R., Callahan, P., Beck, W., Piermarini, E., Terry, A.V., Sullivan, K.A., et al. (2021). A cellular approach to understanding and treating Gulf War Illness. Cell Mol Life Sci 78, 6941–6961. 10.1007/s00018-021-03942-3.

41. Serrao, M., Rinaldi, M., Ranavolo, A., Lacquaniti, F., Martino, G., Leonardi, L., Conte, C., Varrecchia, T., Draicchio, F., Coppola, G., et al. (2016). Gait Patterns in Patients with Hereditary Spastic Paraparesis. PLoS One 11, e0164623. 10.1371/journal.pone.0164623.

42. Parasuraman, S., Raveendran, R., and Kesavan, R. (2010). Blood sample collection in small laboratory animals. J Pharmacol Pharmacother 1, 87–93. 10.4103/0976-500x.72350.

43. Cao, S.Y., Hu, Y., Chen, C., Yuan, F., Xu, M., Li, Q., Fang, K.H., Chen, Y., and Liu, Y. (2017). Enhanced derivation of human pluripotent stem cell-derived cortical glutamatergic neurons by a small molecule. Sci Rep 7, 3282. 10.1038/s41598-017-03519-w.

44. Gaspard, N., Bouschet, T., Herpoel, A., Naeije, G., van den Ameele, J., and Vanderhaeghen, P. (2009). Generation of cortical neurons from mouse embryonic stem cells. Nat Protoc 4, 1454–1463. 10.1038/nprot.2009.157.

45. Arlotta, P., Molyneaux, B.J., Chen, J., Inoue, J., Kominami, R., and Macklis, J.D. (2005). Neuronal subtype-specific genes that control corticospinal motor neuron development in vivo. Neuron 45, 207–221. 10.1016/j.neuron.2004.12.036.

46. Genç, B., Jara, J.H., Sanchez, S.S., Lagrimas, A.K.B., Gözütok, Ö., Koçak, N., Zhu, Y., and Hande Özdinler, P. (2022). Upper motor neurons are a target for gene therapy and UCHL1 is necessary and sufficient to improve cellular integrity of diseased upper motor neurons. Gene Ther 29, 178–192. 10.1038/s41434-021-00303-4.

47. Yasvoina, M.V., Genç, B., Jara, J.H., Sheets, P.L., Quinlan, K.A., Milosevic, A., Shepherd, G.M., Heckman, C.J., and Özdinler, P.H. (2013). eGFP expression under UCHL1 promoter genetically labels corticospinal motor neurons and a subpopulation of degeneration-resistant spinal motor neurons in an ALS mouse model. J Neurosci 33, 7890–7904. 10.1523/jneurosci.2787-12.2013.

48. Day, I.N., and Thompson, R.J. (2010). UCHL1 (PGP 9.5): neuronal biomarker and ubiquitin system protein. Prog Neurobiol 90, 327–362. 10.1016/j.pneurobio.2009.10.020.

49. Osaka, H., Wang, Y.L., Takada, K., Takizawa, S., Setsuie, R., Li, H., Sato, Y., Nishikawa, K., Sun, Y.J., Sakurai, M., et al. (2003). Ubiquitin carboxy-terminal hydrolase L1 binds to and stabilizes monoubiquitin in neuron. Hum Mol Genet 12, 1945–1958. 10.1093/hmg/ddg211.

50. Parodi, L., Fenu, S., Barbier, M., Banneau, G., Duyckaerts, C., Tezenas du Montcel, S., Monin, M.L., Ait Said, S., Guegan, J., Tallaksen, C.M.E., et al. (2018). Spastic paraplegia due to SPAST mutations is modified by the underlying mutation and sex. Brain 141, 3331–3342. 10.1093/brain/awy285.

51. Wang, J., Wu, Y., Dong, H., Ji, Y., Zhang, L., Liu, Y., Liu, Y., Gao, X., Jia, Y., and Wang, X. (2023). A novel truncated variant in SPAST results in spastin accumulation and defects in microtubule dynamics. BMC Med Genomics 16, 321. 10.1186/s12920-023-01759-6.

52. Errico, A., Ballabio, A., and Rugarli, E.I. (2002). Spastin, the protein mutated in autosomal dominant hereditary spastic paraplegia, is involved in microtubule dynamics. Hum Mol Genet 11, 153–163. 10.1093/hmg/11.2.153.

53. Yandamuri, S.S., and Lane, T.E. (2016). Imaging Axonal Degeneration and Repair in Preclinical Animal Models of Multiple Sclerosis. Front Immunol 7, 189. 10.3389/fimmu.2016.00189.

54. Tsyba, D.L., Kirik, O.V., and Korzhevskii, D.E. (2021). SMI-32 — a novel axonal injury marker for investigation of ischemic brain pathology. MAJ 20, 63–68. 10.17816/maj49849.

55. McIlwain, D.L., and Hoke, V.B. (2005). The role of the cytoskeleton in cell body enlargement, increased nuclear eccentricity and chromatolysis in axotomized spinal motor neurons. BMC Neurosci 6, 19. 10.1186/1471-2202-6-19.

56. Leonhardt, M., Sellmer, A., Krämer, O.H., Dove, S., Elz, S., Kraus, B., Beyer, M., and Mahboobi, S. (2018). Design and biological evaluation of tetrahydro-β-carboline derivatives as highly potent histone deacetylase 6 (HDAC6) inhibitors. Eur J Med Chem 152, 329–357. 10.1016/j.ejmech.2018.04.046.

57. Li, Z.Y., Zhang, C., Zhang, Y., Chen, L., Chen, B.D., Li, Q.Z., Zhang, X.J., and Li, W.P. (2017). A novel HDAC6 inhibitor Tubastatin A: Controls HDAC6-p97/VCP-mediated ubiquitination-autophagy turnover and reverses Temozolomide-induced ER stress-tolerance in GBM cells. Cancer Lett 391, 89–99. 10.1016/j.canlet.2017.01.025.

58. Shen, Z., Ji, K., Cai, Z., Huang, C., He, X., Xu, H., and Chen, G. (2021). Inhibition of HDAC6 by Tubastatin A reduces chondrocyte oxidative stress in chondrocytes and ameliorates mouse osteoarthritis by activating autophagy. Aging (Albany NY) 13, 9820–9837. 10.18632/aging.202736.

59. Rehbach, K., Kesavan, J., Hauser, S., Ritzenhofen, S., Jungverdorben, J., Schüle, R., Schöls, L., Peitz, M., and Brüstle, O. (2019). Multiparametric rapid screening of neuronal process pathology for drug target identification in HSP patient-specific neurons. Sci Rep 9, 9615. 10.1038/s41598-019-45246-4.

60. Navas-Sánchez, F.J., Martín De Blas, D., Fernández-Pena, A., Alemán-Gómez, Y., Lage-Castellanos, A., Marcos-Vidal, L., Guzmán-De-Villoria, J.A., Catalina, I., Lillo, L., Muñoz-Blanco, J.L., et al. (2022). Corticospinal tract and motor cortex degeneration in pure hereditary spastic paraparesis type 4 (SPG4). Amyotroph Lateral Scler Frontotemporal Degener 23, 25–34. 10.1080/21678421.2021.1962353.

61. McDermott, C.J., Burness, C.E., Kirby, J., Cox, L.E., Rao, D.G., Hewamadduma, C., Sharrack, B., Hadjivassiliou, M., Chinnery, P.F., Dalton, A., and Shaw, P.J. (2006). Clinical features of hereditary spastic paraplegia due to spastin mutation. Neurology 67, 45–51. 10.1212/01.wnl.0000223315.62404.00.

62. Plaud, C., Joshi, V., Kajevu, N., Poüs, C., Curmi, P.A., and Burgo, A. (2018). Functional differences of short and long isoforms of spastin harboring missense mutation. Dis Model Mech 11. 10.1242/dmm.033704.

63. Solowska, J.M., Morfini, G., Falnikar, A., Himes, B.T., Brady, S.T., Huang, D., and Baas, P.W. (2008). Quantitative and functional analyses of spastin in the nervous system: implications for hereditary spastic paraplegia. J Neurosci 28, 2147–2157. 10.1523/jneurosci.3159-07.2008.

64. Denton, K.R., Lei, L., Grenier, J., Rodionov, V., Blackstone, C., and Li, X.J. (2014). Loss of spastin function results in disease-specific axonal defects in human pluripotent stem cell-based models of hereditary spastic paraplegia. Stem Cells 32, 414–423. 10.1002/stem.1569.

65. Trotta, N., Orso, G., Rossetto, M.G., Daga, A., and Broadie, K. (2004). The hereditary spastic paraplegia gene, spastin, regulates microtubule stability to modulate synaptic structure and function. Curr Biol 14, 1135–1147. 10.1016/j.cub.2004.06.058.

66. Fassier, C., Tarrade, A., Peris, L., Courageot, S., Mailly, P., Dalard, C., Delga, S., Roblot, N., Lefèvre, J., Job, D., et al. (2013). Microtubule-targeting drugs rescue axonal swellings in cortical neurons from spastin knockout mice. Dis Model Mech 6, 72–83. 10.1242/dmm.008946.

67. Abrahamsen, G., Fan, Y., Matigian, N., Wali, G., Bellette, B., Sutharsan, R., Raju, J., Wood, S.A., Veivers, D., Sue, C.M., and Mackay-Sim, A. (2013). A patient-derived stem cell model of hereditary spastic paraplegia with SPAST mutations. Dis Model Mech 6, 489–502. 10.1242/dmm.010884.

