## Supplementary table for "Genotype–Phenotype Distinctions in Spastic Paraplegia 4 Reveal HDAC6 as a Therapeutic Target"

### Supplementary Tables

Supplementary Table 1: List of antibodies, primers, and plasmids used for this study.

| Reagent or Resource | Source | Catalog Number |
|---|---|---|
| **Antibodies for IHC and WB** | | |
| SOX2 | R&D Systems | AF2018 |
| Neun | Abcam | ab104224 |
| MAP2 | Novus Biologicals | NB300-213 |
| βIII- Tubulin | BioLegend | 801201 |
| βIII- Tubulin | BioLegend | 802001 |
| CTIP2 | Abcam | ab18465 |
| GFAP | Abcam | ab53554 |
| SMI312 | BioLegend | 837904 |
| L1CAM | BioLegend | 371607 |
| vGlut1 | Synaptic Systems | 135302 |
| Synaptophysin | Proteintech | 60191-1-AP |
| SMI32 | BioLegend | 801701 |
| Acetylated Tubulin | Sigma-Aldrich | T6793 |
| HDAC6 | Cell Signaling Technology | 7612S |
| βIII- Tubulin | Abcam | ab7751 |
| GAPDH | Abcam | ab8245 |
| Spastin | Abcam | ab77144 |
| Anti-Flag | Proteintech | 20543-1-AP |
| Anti-Phosphoserine/threonine | Amyjet Scientific | PPS-PP2551 |
| Anti-HDAC6 | Affinity Biosciences | AF6485 |
| Anti-mCherry | Abcam | ab205402 |
| **Primers for RT-qPCR** | | |
| CTIP2<br>FWD: ATGTCCCGCCGCAAACA<br>REV: TCGTCTTCTTCGAGGATGG | Integrated DNA Technologies, Inc. | N/A |
| L1CAM<br>FWD: GTCCACTACACCTTTAGGGTTAC<br>REV: CTTCACATCCACAGGGTTCTT | Integrated DNA Technologies, Inc. | N/A |
| CRYM<br>FWD: GAGCTGGGAGAAGTGATTAAGG<br>REV: TGGCTGCAACTGTGTCTT | Integrated DNA Technologies, Inc. | N/A |
| vGlut1<br>FWD: CACTCAGATTCCAGGAGGATTT<br>REV: CGAAGATGACACAGCCATAGT | Integrated DNA Technologies, Inc. | N/A |
| UCLH1<br>FWD: CCAATGTCGGGTAGATGACAAG<br>REV: AGGCATTCGTCCATCAAGTTC | Integrated DNA Technologies, Inc. | N/A |
| N-terminal Spastin<br>FWD: CAGAAGGAGCAAGCTGTGGA<br>REV: GTAAGCGGTCCTTGGCCATA | Integrated DNA Technologies, Inc. | N/A |
| C-terminal Spastin FWD: AGAAGAGAAGGGGAGCACGA<br>REV: AAACGCCTGAGAACAGCCTC | Integrated DNA Technologies, Inc. | N/A |
| GAPDH<br>FWD: TCGGAGTCAACGGATTTGGT<br>REV: TTCCCGTTCTCAGCCTTGAC | Integrated DNA Technologies, Inc. | N/A |
| **Plasmids for overexpression studies and co-IP** | | |
| Wildtype M87 | Addgene | 89322 |
| M87 184X | Addgene | 92367 |

| | | |
|---|---|---|
| M87 C448Y | Addgene | 89320 |
| M87 S245X | Addgene | 92368 |
| Wildtype M1 | Addgene | 89321 |
| M1 184X | Addgene | 92365 |
| M1 C448Y | Addgene | 89319 |
| M1 S245X | Addgene | 92366 |
| M1 L195V-Myc tagged | Addgene | 87716 |
| M1 E112K-Myc tagged | Addgene | 87715 |
| mcherry-HDAC6 | Generated in Lab | N/A |
| CON255-Flag | Generated in Lab | N/A |
| M1-Spastin$^{WT}$-Flag | Generated in Lab | N/A |
| M87-Spastin$^{WT}$-Flag | Generated in Lab | N/A |
| M1-Spastin$^{C448Y}$-Flag | Generated in Lab | N/A |
| M1-Spastin$^{S245X}$-Flag | Generated in Lab | N/A |

Supplementary Table 2: Mean ± SD values from Figure 1.

| | **Cortical Organoids** | **Motor Cortical Organoids** |
|---|---|---|
| Relative gene expression- CTIP2 | 1.013 ± 0.202 | 2.077 ± 0.040 |
| Relative gene expression- L1CAM | 1.003 ± 0.099 | 5.631 ± 0.905 |
| Relative gene expression- vGlut1 | 1.003 ± 0.091 | 5.733 ± 1.658 |
| Relative gene expression- CRYM1 | 1.006 ± 0.137 | 5.105 ± 1.212 |
| Relative gene expression- UCLH1 | 1.007 ± 0.144 | 1.004 ± 0.067 |

Supplementary Table 3: Mean ± SD values from Figure 2.

| | SPAST$^{WT/WT}$ | SPAST$^{WT/S245X}$ | SPAST$^{WT/C448Y}$ |
|---|---|---|---|
| Relative gene expression (N- Terminal Primers) | 1.006 ± 0.133 | 5.103 ± 2.456 | 7.266 ± 0.959 |
| Relative gene expression-C- Terminal Primers | 1.023 ± 0.253 | 3.303 ± 0.582 | 2.480 ± 0.443 |
| Full length M1-Spastin/GAPDH | 1.000 ± 0.000 | 0.879 ± 0.213 | 1.463 ± 0.206 |
| Full length M87-Spastin/GAPDH | 1.000 ± 0.000 | 0.691 ± 0.094 | 1.238 ± 0.132 |
| 1M ratio of SMI32/ΒIII-tubulin | 1.000 ± 0.619 | 0.651 ± 0.396 | 1.260 ± 1.257 |
| 3M ratio of SMI32/ΒIII-tubulin | 1.000 ± 0.266 | 1.169 ± 0.277 | 1.664 ± 0.516 |
| 6M ratio of SMI32/ΒIII-tubulin | 1.112 ± 0.530 | 2.459 ± 1.263 | 4.340 ± 1.924 |
| No. of CTIP2+ cells/100μm$^2$ | 0.153 ± 0.019 | 0.135 ± 0.012 | 0.023 ± 0.008 |
| CTIP2+ cell body perimeter (μm) | 0.562 ± 0.120 | 0.634 ± 0.112 | 1.118 ± 0.107 |
| 3M Mean Spike Count in 5mins | 44424.250 ± 4833.034 | 29455.047 ± 2666.436 | 70030.380 ± 4793.515 |
| 6M Mean Spike Count in 5mins | 36197.333 ± 6851.016 | 52181.517 ± 2690.224 | 11250.484 ± 3918.244 |
| 3M Mean Spike Rate in 5mins (Hz) | 166.858 ± 11.772 | 84.864 ± 5.114 | 233.035 ± 20.090 |
| 6M Mean Spike Rate in 5mins (Hz) | 144.427 ± 5.998 | 165.624 ± 6.958 | 46.095 ± 5.671 |
| 3M Mean Burst Duration in 5mins (s) | 291584.000 ± 20000.000 | 163513.133 ± 12377.599 | 338387.233 ± 38811.773 |
| 6M Mean Burst Duration in 5mins (s) | 253899.633 ± 15676.727 | 88020.530 ± 37754.360 | 4739.680 ± 1907.169 |
| 3M Mean Spikes in Bursts in 5mins | 39058.693 ± 8516.771 | 15727.660 ± 3292.816 | 71399.507 ± 2794.982 |
| 6M Mean Spikes in Bursts in 5mins | 33380.277 ± 3476.733 | 9825.749 ± 3611.799 | 1088.240 ± 823.148 |
| 3M Mean Network Burst Count in 5mins | 2675349.667 ± 198205.349 | 1644156.333 ± 149237.050 | 4168442.000 ± 287501.941 |
| 6M Mean Network Burst Count in 5mins | 2296725.667 ± 240139.396 | 1210889.000 ± 267244.142 | 675027.667 ± 235095.121 |
| 3M Mean Network Burst Frequency (Hz) | 152.778 ± 10.199 | 93.658 ± 5.202 | 249.459 ± 10.243 |
| 6M Mean Network Burst Frequency (Hz) | 157.457 ± 19.008 | 67.244 ± 14.851 | 37.479 ± 13.048 |

Supplementary Table 4: Mean ± SD values from Figure 3.

| | SPAST$^{WT/WT}$ | SPAST$^{WT/WT}$ + Tub A | SPAST$^{WT/S245X}$ | SPAST$^{WT/S245X}$ + Tub A | SPAST$^{WT/C448Y}$ | SPAST$^{WT/C448Y}$ + Tub A |
|---|---|---|---|---|---|---|
| 3M ratio of Acetylated-tubulin/ΒIII-tubulin | 1.000 ± 0.145 | 1.146 ± 0.151 | 1.131 ± 0.315 | 0.929 ± 0.034 | 0.709 ± 0.056 | 1.171 ± 0.049 |
| 6M ratio of Acetylated-tubulin/ΒIII-tubulin | 1.000 ± 0.029 | 1.030 ± 0.056 | 0.761 ± 0.035 | 1.013 ± 0.042 | 0.754 ± 0.053 | 0.950 ± 0.048 |
| 3M ratio of SMI32/ΒIII-tubulin | 1.000 ± 0.266 | 0.799 ± 0.623 | 1.169 ± 0.277 | 1.468 ± 0.097 | 1.664 ± 0.516 | 1.248 ± 0.188 |
| 6M ratio of SMI32/ΒIII-tubulin | 0.714 ± 0.470 | 2.493 ± 0.318 | 0.957 ± 1.026 | 2.589 ± 0.233 | 0.714 ± 0.867 | 0.714 ± 0.318 |
| 3M HDAC6 Activity (U/mg) | 478.043 ± 24.184 | 85.444 ± 1.007 | 427.472 ± 6.715 | 171.386 ± 6.660 | 1034.397 ± 86.636 | 174.062 ± 4.750 |
| 3M HDAC6 Activity (U/mg) | 115.883 ± 4.037 | 6.619 ± 1.480 | 268.192 ± 12.159 | 21.475 ± 0.279 | 272.817 ± 18.956 | 25.621 ± 1.012 |

Supplementary Table 5: Mean ± SD values from Figure 3.

| | **SPAST$^{WT/WT}$** | **SPAST$^{WT/S245X}$** | **SPAST$^{WT/C448Y}$** |
|---|---|---|---|
| HDAC6/GAPDH | 1.000 ± 0.000 | 0.843 ± 0.451 | 0.840 ± 0.248 |

Supplementary Table 6: Mean ± SD values from Figure 4.

| | **M1-Spastin$^{WT}$-Flag** | **M87-Spastin$^{WT}$-Flag** | **M1-Spastin$^{C448Y}$-Flag** | **M1-Spastin$^{S245X}$-Flag** |
|---|---|---|---|---|
| Relative SPAST protein level (IP:mCherry-HDAC6) | 1.000 ± 0.000 | 0.239 ± 0.103 | 3.639 ± 0.676 | 2.898 ± 0.360 |
| Relative HDAC6 protein level (IP: Flag-SPAST) | 1.000 ± 0.000 | 0.369 ± 0.179 | 3.310 ± 0.543 | 0.867 ± 0.814 |

Supplementary Table 7: Mean ± SD values from Figure 4.

| | **Control** | **M87-WT** | **M87-184X** | **M87-C448Y** | **M87-S245X** | |
|---|---|---|---|---|---|---|
| HDAC6 Activity (U/mg) | 592.671 ± 12.975 | 632.581 ± 43.144 | 572.391 ± 371.650 | 691.071 ± 106.532 | 738.284 ± 87.516 | |
| | **M1-WT** | **M1-184X** | **M1-C448Y** | **M1-S245X** | **M1-L195V** | **M1-E112K** |
| HDAC6 Activity (U/mg) | 659.075 ± 25.054 | 876.999 ± 26.280 | 2068.461 ± 296.268 | 1679.451 ± 167.854 | 1670.676 ± 100.310 | 3407.728 ± 345.665 |

Supplementary Table 8: Chemistry panel of serum extracted from whole blood samples to evaluate hepatotoxicity with Tubastatin A (Tub A) treatment.

| | **Albumin** | **Alkaline Phosphatase** | **Alanine Transaminase** | **Aspartate Aminotransferase** |
|---|---|---|---|---|
| Reference Range (Units) | 2.72 - 4.2 (g/dL) | 65.5 - 364.2 (U/L) | 24.3 - 114.25 (U/L) | 39.55 - 386.05 (U/L) |
| WT + Veh | 4.37 ± 0.42 | 35.67 ± 11.59 | 71.33 ± 11.37 | 352.33 ± 191.81 |
| WT + Tub A | 6.67 ± 1.21 | 35.33 ± 2.52 | 38.00 ± 47.29 | 147.00 ± 15.52 |
| dHet + Veh | 6.63 ± 1.55 | 70.33 ± 6.11 | 76.00 ± 11.36 | 509.67 ± 26.10 |
| dHet + Tub A | 5.40 ± 1.11 | 50.67 ± 3.51 | 94.67 ± 25.38 | 119.33 ± 9.50 |

Data shown as mean ± SD. n=3 mice per group.

Supplementary Table 9: Mean ± SD values from Figure 5.

| | **WT + Veh** | **WT + Tub A** | **dHet + Veh** | **dHet + Tub A** |
|---|---|---|---|---|
| Print Length | 1.330 ± 0.193 | 1.328 ± 0.182 | 1.159 ± 0.140 | 1.263 ± 0.112 |
| Print Width | 1.266 ± 0.188 | 1.261 ± 0.127 | 1.114 ± 0.173 | 1.224 ± 0.126 |
| Print Area | 1.234 ± 0.326 | 1.147 ± 0.218 | 0.897 ± 0.203 | 1.076 ± 0.184 |
| Print Position | 1.214 ± 0.368 | 1.105 ± 0.500 | 0.964 ± 0.528 | 1.221 ± 0.507 |
| HDAC6 Activity | 158.421 ± 13.901 | 137.349 ± 6.334 | 207.235 ± 8.971 | 117.410 ± 16.201 |
| Acetylated-tubulin/BIII-tubulin | 0.884 ± 0.023 | 0.885 ± 0.069 | 0.622 ± 0.056 | 0.946 ± 0.052 |
| BIII-tubulin/GAPDH | 4.225 ± 0.685 | 4.181 ± 0.911 | 4.050 ± 0.687 | 4.333 ± 0.577 |
| Total Number of Axons | 0.424 ± 0.167 | 0.366 ± 0.116 | 0.178 ± 0.113 | 0.253 ± 0.067 |
| Average Axonal Perimeter | 4.264 ± 2.284 | 4.653 ± 3.240 | 5.088 ± 3.599 | 4.156 ± 2.244 |
| Total Number of Axons-Normal | 0.230 ± 0.108 | 0.240 ± 0.080 | 0.159 ± 0.086 | 0.207 ± 0.059 |
| Total Number of Axons-Swollen | 0.078 ± 0.034 | 0.059 ± 0.027 | 0.157 ± 0.073 | 0.126 ± 0.078 |
